# Over-accessible chromatin links myeloma initiating genetic events to oncogenic transcriptomes and aberrant transcription factor regulatory networks

**DOI:** 10.1101/2020.06.11.140855

**Authors:** Jaime Alvarez-Benayas, Alexia Katsarou, Nikolaos Trasanidis, Aristeidis Chaidos, Philippa C May, Kanagaraju Ponnusamy, Xiaolin Xiao, Marco Bua, Maria Atta, Irene AG Roberts, Holger W Auner, Evdoxia Hatjiharissi, Maria Papaioannou M, Valentina S Caputo, Ian M Sudbery IM, Anastasios Karadimitris

**Affiliations:** Department of Molecular Biology and Biotechnology, University of Sheffield, Sheffield, United Kingdom; Hugh & Josseline Langmuir Centre for Myeloma Research, Centre for Haematology, Department of Immunology and Inflammation, Imperial College London, London, United Kingdom; Department of Haematology, Hammersmith Hospital, Imperial College Healthcare NHS Foundation Trust, London, United Kingdom; MRC Molecular Haematology Unit and Paediatrics, MRC Weatherall Institute of Molecular Medicine, Oxford, United Kingdom; First Department of Internal Medicine, Division of Haematology, AHEPA University Hospital, Aristotle University of Thessaloniki, Thessaloniki, Greece

## Abstract

Multiple myeloma is a genetically heterogeneous cancer of the bone marrow plasma cells (PC). Myeloma initiating genetic events define subgroups (MIE) and drive distinct oncogenic transcriptomes that converge into a mutually exclusive overexpression of *CCND1* and *CCND2* oncogenes. Here, with reference to normal PC, we dissect how MIE impact the chromatin regulatory landscape of MM. We find that chromatin accessibility combined with transcriptome profiling classifies myeloma genetic subgroups, while in a topologically constrained manner, distal rather than proximal regulatory elements influence myeloma transcriptomes. Across and within MIE-defined subgroups, genes and pathways critical for myeloma biology can be linked to developmentally activated or *de novo* formed enhancers. We show that existing transcription factors, co-opted to organise highly ordered, aberrant regulatory networks, generate known and novel myeloma cell dependencies and help identify prognostic markers. Finally, we discover and functionally validate the critical enhancer that regulates ectopic expression of *CCND2* in MM.

---

MM is a common, genetically heterogeneous incurable cancer of the bone marrow plasma cells (PC), the terminally differentiated immunoglobulin-secreting B lineage cells (Palumbo and Anderson, 2011). Distinct transcriptome profiles in MM reflect two categories of myeloma initiating genetic events (MIE): Over-expression in up to half of cases of oncogenes such as *CCND1, MAF* and *MMSET* by their juxtaposition to the IgH enhancer in the t(11;14), t(14;16) and t(4;14) cytogenetic subgroups respectively; hyperdiploidy (HD) is the MIE driving oncogenic transcriptomes in the rest of cases (Manier et al., 2017; Morgan et al., 2012). High frequency secondary copy number aberrations, single nucleotide variants and indels further shape the distinct impact of the MIE and generate extensive genetic heterogeneity (Chapman et al., 2011; Lohr et al., 2014; Walker et al., 2015). This heterogeneity converges, in most cases, to a functionally dichotomous, mutually exclusive overexpression of the cell cycle regulators CCND1 and CCND2 to which myeloma PC remain addicted, irrespective of primary or secondary genetic events (Bergsagel et al., 2005; Ely et al., 2005; Tiedemann et al., 2008).

As well as over-expression by juxtaposition to the IgH enhancer in 20% of MM, *CCND1* over-expression is associated with chr11q25 gain in over half of HD cases (Shah et al., 2018). However, the transcriptional mechanisms that result in *CCND2* over-expression, seen in nearly 50% of MM cases and spanning all genetic subgroups except t(11;14), are not known.

Chromatin accessibility profiling by ATAC-seq has been used to characterise the regulatory landscape of hundreds of different solid tumour cancers and in combination with other datatypes, has demonstrated its utility in categorising cancer and in the discovery of distal regulatory elements such as candidate enhancers of critical oncogenes (Corces et al., 2018). Further, by means of transcription factor (TF) footprinting, ATAC-seq allows inference of TF binding profiles and construction of relevant regulatory gene networks or circuitries (Ott et al., 2018; Rendeiro et al., 2016). Such networks may help identify TF with previously unrecognised roles in the biology of a given cancer. The power of discovery of such approaches is enhanced by comparing chromatin accessibility of the cancer of interest with that of the cell(s) of origin; however, these in many cases are unknown or uncertain. Indeed, one of the limitations of the pan-cancer chromatin accessibility maps was lack of normal cell-of-origin-derived chromatin accessibility profiles as reference (Corces et al., 2018).

In MM, a previous study of chromatin accessibility, using as reference PC derived *in vitro* from mature splenic B cells, demonstrated that chromatin alterations, including heterochromatin de-condensation, underpin pan-myeloma regulatory gene networks that involve known and novel TF such as FLI1 (Jin et al., 2018). However, how myeloma initiating events are linked to chromatin accessibility and chromatin regulatory status; how chromatin changes impact the transcriptomes and biological pathways associated with each myeloma initiating event and how these compare with reference to normal BM PC i.e., the cells of origin of MM and to cells of the B cell developmental trajectory, have not been addressed.

Here we show that by using chromatin accessibility as an integrating platform of multiple datatypes we can link MIE to their oncogenic transcriptomes and biological pathways. Through this process we discover novel, *in cis* and *in trans*, regulators of myeloma biology, including those involved in the regulation and aberrant expression of *CCND2* in MM.

## Results

### ATAC-seq reveals an increased chromatin accessibility in myeloma PC and helps classify genetic subgroups

To link MIE with chromatin and transcriptional changes we obtained fresh, highly purified BM (CD19^+/-^) PC from 3 healthy normal donors (ND) and myeloma PC from 30 MM patients, covering the main MIE subgroups as defined by fluorescent in situ hybridization **(Figure 1a, Suppl Fig 1a and Suppl Table 1).** For each sample we obtained paired chromatin accessibility and transcriptome profiles by ATAC-seq and RNA-seq respectively.

**Figure 1.**
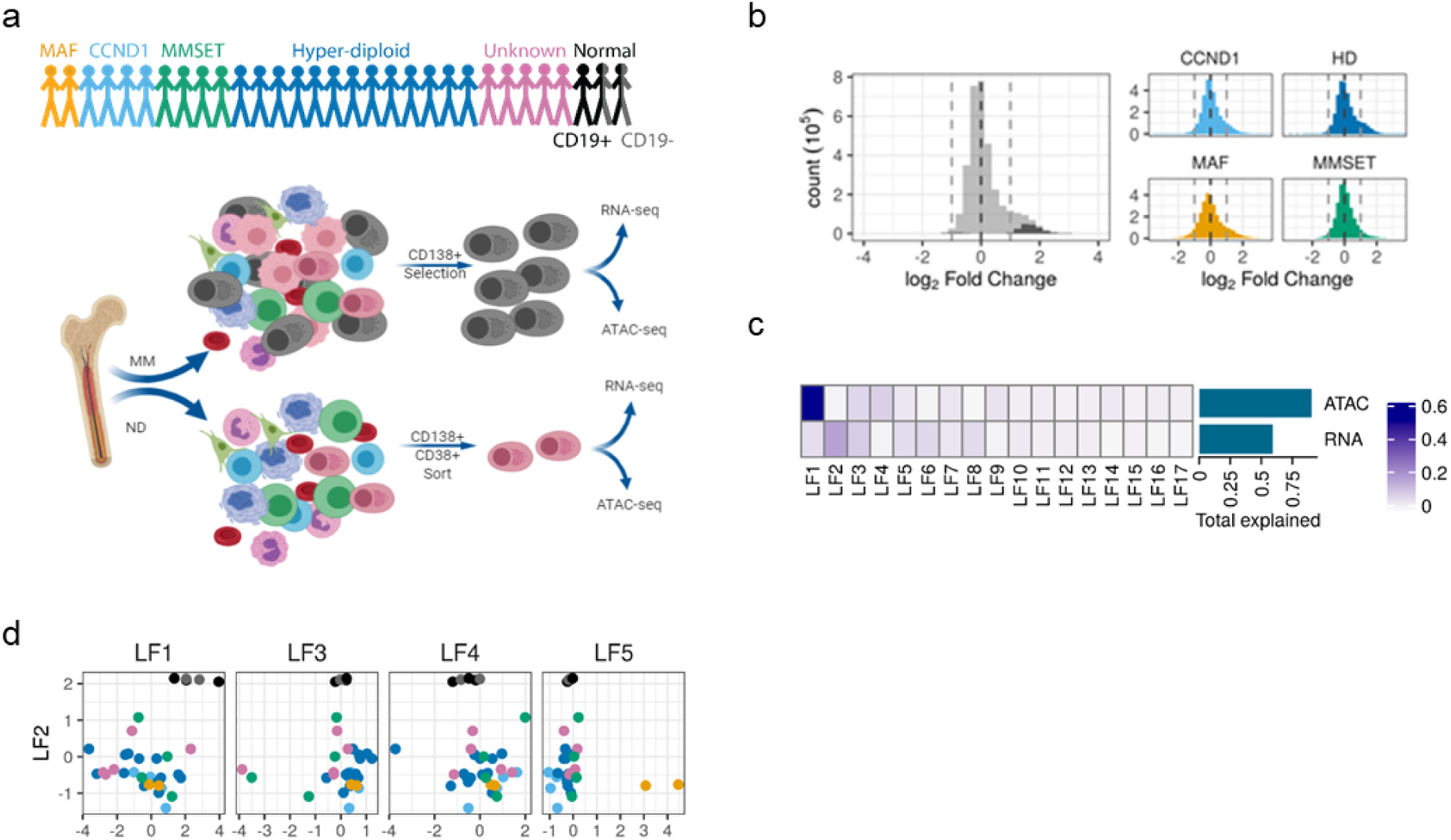
Over-accessible distal chromatin classifies myeloma genetic subgroups. **a)** Study patient population and design. Transcriptome and chromatin accessibility were assessed by RNA-seq and ATAC-seq respectively in myeloma PC from 30 MM patients and PC from 3 healthy donors. For two of the controls we obtained samples of both CD19^+^ and CD19^-^ PC. **b)** Change in average ATAC signal over pan-myeloma (left) and subgroup (right) peaks expressed as normalized log_2_ myeloma/normal PC read count. **c)** Variance explained by the first 17 latent factors (LF) for each of RNA and ATAC signal as calculated by Multi-Omics Factor Analysis (MOFA). **d)** LF scores for each sample for the first 5 latent factors in the MOFA model. Subtype for each sample is denoted by colour, as per a). LF1 and LF2 distinguish normal from MM samples, LF5 distinguishes MAF, CCND1 and MMSET samples from the rest.

ATAC-seq generated 295,238 peaks from all samples signifying chromatin accessibility **(Suppl Fig 1b)**. These peaks were non-randomly distributed on promoters, coding and intergenic genomics regions and were either shared by samples of two or more genetic subgroups or were present only in samples of one subgroup **(Suppl Fig 1b & c**). In each genetic subgroup and overall, chromatin accessibility of myeloma PC was enhanced in comparison to normal PC **(Figure 1b).** Transcriptome profiling **(Suppl Fig 1d)** identified 3,036 differentially expressed genes (DEG) between myeloma subgroups and ND PC, including overexpression of known MIE-driven oncogenes in myeloma PC and highlighted the mutually exclusive expression of *CCND1* vs *CCND2* **(Suppl Fig 1d)**.

By employing the machine learning Multi-Omics Factor Analysis (MOFA) approach(Argelaguet et al., 2018), we explored in an unsupervised manner how differences in chromatin accessibility and transcriptome might help classify MM genetic subgroups **(Figure 1c & d, Suppl Fig 1e).** We found that despite the range in means being 20-fold higher for expression, in the combined accessibility-transcriptome model, altered chromatin accessibility accounted for more variance than expression **(Figure 1c)**. Of the top five identified MOFA latent factors (LF), the mostly chromatin accessibility-driven LF1 and the mainly transcriptome-driven LF2 distinguished ND from myeloma PC, whilst two more separated *MMSET, MAF* and *CCND1* subgroups **(Figure 1d).** Further, the combination of datatypes provided a clearer separation of subgroups than either alone **(Suppl Fig 1f&g).** Since there was no difference between CD19^+^ and CD19^-^ ND PC cells, they were merged for subsequent analyses.

Thus, over-accessible chromatin provides a framework for additional functional categorisation of myeloma genetic subgroups.

### Distal rather than proximal elements regulate myeloma oncogenic transcriptomes

To assess the functional impact of over-accessible chromatin in myeloma PC, with reference to ND PC, we correlated accessibility with transcriptome changes. This showed that in myeloma PC, genes with more open peaks at their transcription start sites (TSS) were only slightly more likely to be over-expressed than those without (3.5% vs 2.7% OR=1.3, p=0.01). Meanwhile, genes with a distal, non-TSS overlapping, significantly more accessible peak within 500kb from the TSS were more likely to be over-expressed (3.5% vs 2.0%, OR=1.7, p=1.5×10^−8^). In addition, over-expressed genes were associated with significantly more such peaks than non-overexpressed genes (**Figure 2a**), and the distance from over-expressed genes to the closest such peak was shorter (**Figure 2b**).

**Figure 2.**
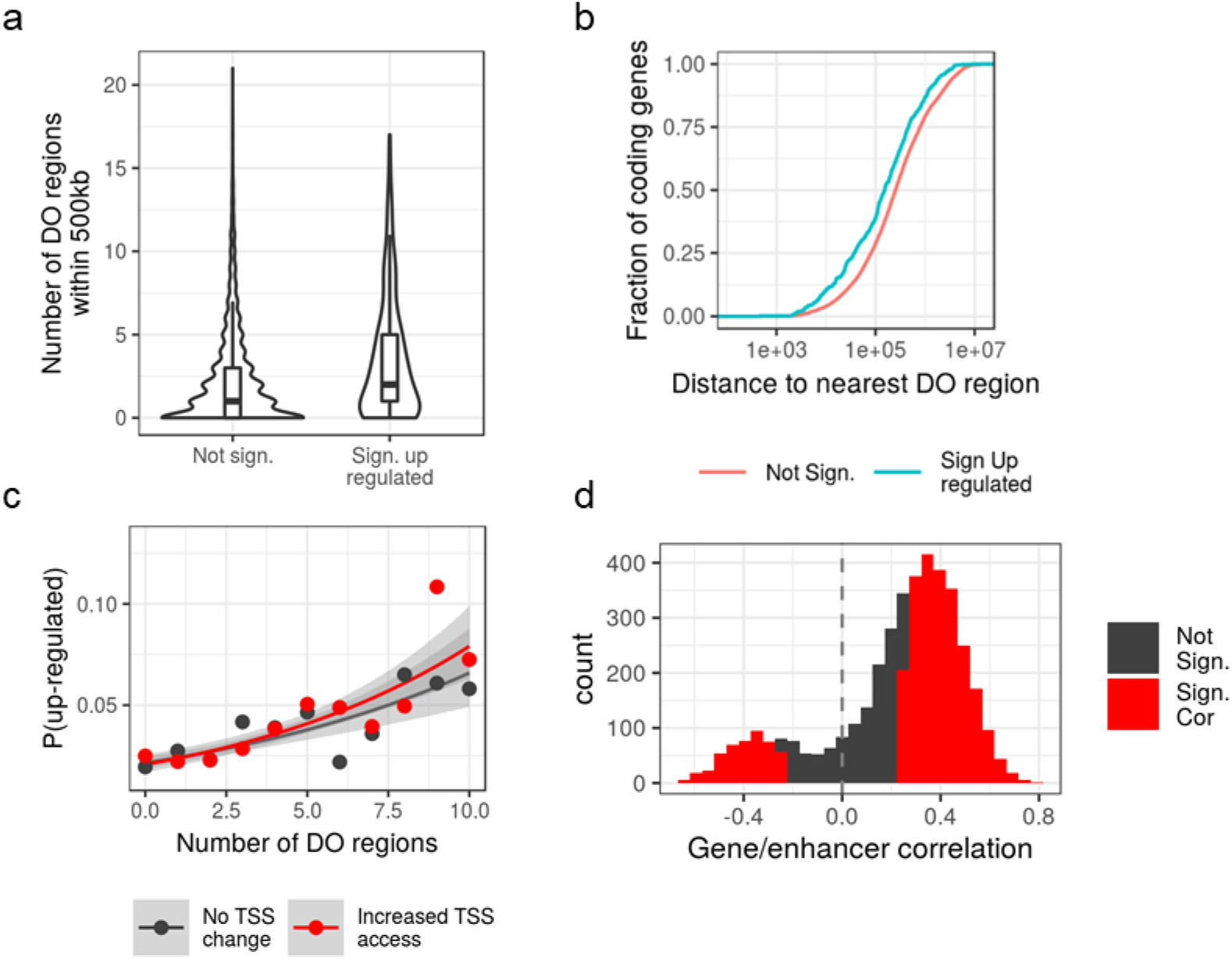
Role of distal regulatory elements in the regulation of myeloma transcriptomes. **a)** Number of pan-myeloma peaks found to be significantly differentially open (DO) in myeloma samples within 500kb of genes found to be significantly upregulated in myeloma (Sign. up regulated), or not (Not sign.). **b)** Cumulative distribution function for the distance to the TSS of the nearest pan-myeloma peak found to be significantly more open in myeloma for genes that either significantly upregulated in myeloma (Sign. Up regulated, blue) or not (Not sign., red). **c)** Logistic regression of number of DO regions within 500kb of a gene (x-axis), whether a gene has a significantly more open promoter in myeloma (red – yes, black – no) on the probability of the gene being differentially upregulated in myeloma. Points represent fraction of genes up-regulated with a given number of DO regions within 500kb and TSS status, and lines represent model fit, grey ribbon 95% confidence interval. Genes with 10 or more DO regions were pooled together. **d)** Correlation of signal for differentially accessible ATAC regions and differentially expressed genes within the same topology associating domain (TAD).

Similarly, in a combined logistic regression analysis, number of more accessible non-TSS regions was predictive of over-expression (p<2×10^−16^) but a more open promoter was not (p=0.5**; Figure 2c)**.

In addition, using as reference the 3D genome architecture of EBV transformed B cells(Rao et al., 2014), the same lineage as PC, within the same topologically associating domain (TAD), we found a significant enrichment for differentially accessible regions correlating with differentially expressed genes (Okonechnikov et al., 2019) **(Figure 2d)** further supporting their regulatory association.

Thus, within the topological limits of TADs, chromatin accessibility changes at distal, non-TSS overlapping regions (i.e., candidate enhancers) rather than at promoters are associated with gene over-expression in the cancer state.

### The myeloma ‘enhanceome’ regulates known and novel genes and biological pathways

These distal genomic elements with the higher regulatory potential are likely to represent candidate enhancers and together comprise the myeloma ‘enhanceome’. In total, we identified 4,635 such non-TSS genomic regions with differential accessibility between at least one MM subgroup and ND PC. We then concentrated on the 4,199 differentially accessible regions-DEG pairs that are predicted to be associated within 1Mb of one another. These correspond to 2,581 unique differentially accessible regions (603 in CCND1, 646 in HD, 1,792 in MAF and 873 in MMSET) and clustering of their accessibility signal separated the different myeloma subgroups from each other **(Figure 3a).** While this provided further evidence that distinct chromatin accessibility changes are shaped by MIE it also highlighted regions that overlap between different myeloma subgroups. The MAF subgroup demonstrated the highest number of differentially accessible distal regions, some of which were unique to MAF while others were shared with the other genetic subgroups.

**Figure 3.**
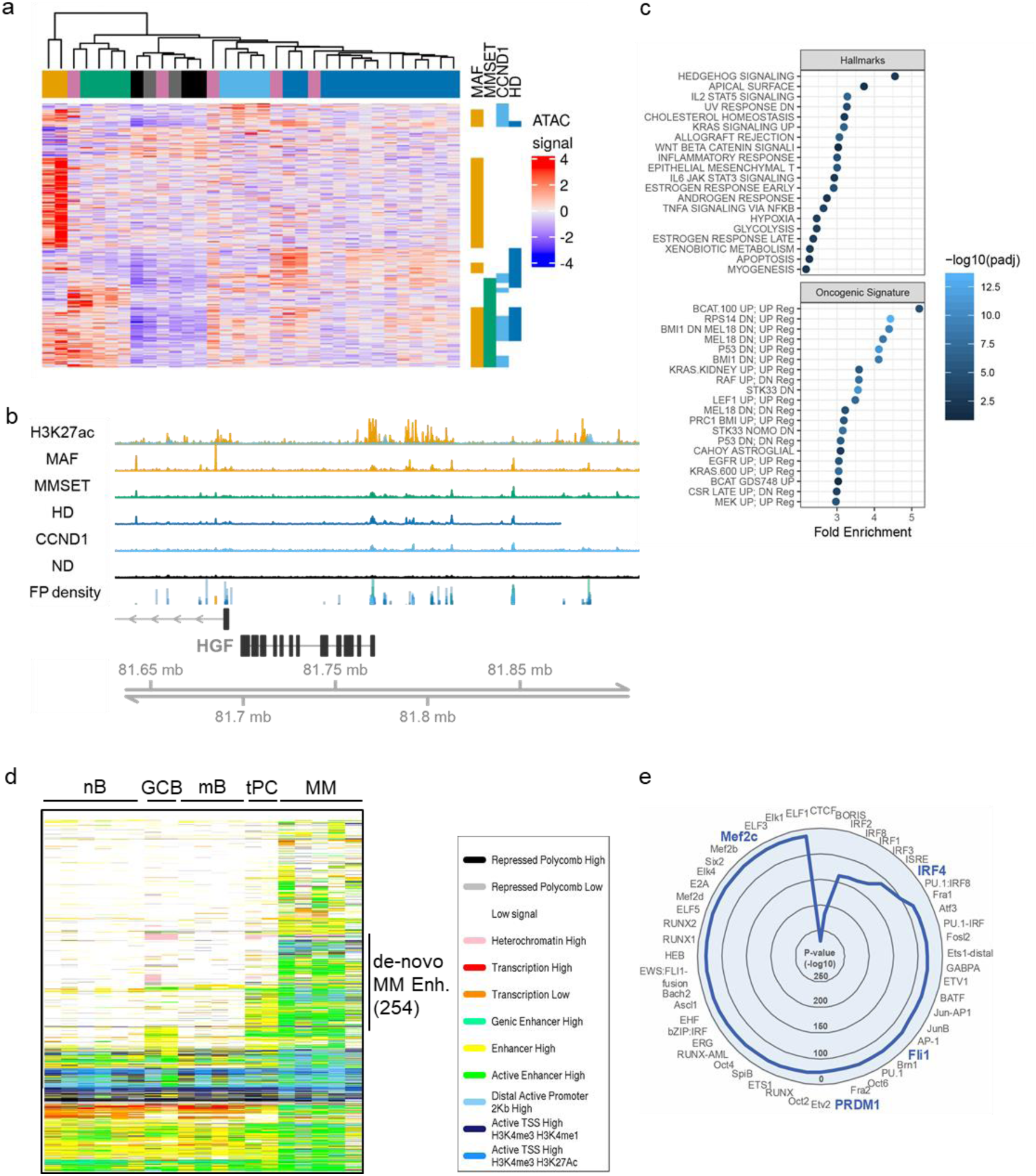
Developmental origins and oncogenic pathways regulated by the myeloma ‘enhanceome’. **a)** Heatmap showing the ATAC signal for all peaks found to be differentially open and within 1Mb of a significantly differentially regulated gene. Data is row scaled. Samples are clustered using Pearson’s correlation distance and the bar above the columns shows the subtype of the sample coded as in Figure 1a, i.e., MAF: orange, CCND1: light blue, MMSET: green, HD: blue, others: pink and ND PC: black/grey. Vertical b ars to the right highlight regions where signal is >2-fold different to ND PC. **b)** Example region around *HGF* showing: average ATAC-seq signal in each subtype; H3K27ac signal in a *MAF*-translocated cell line (orange) or *CCND1*-translocated cell line (blue); FP density: density of footprints in each of the ATAC-seq signals. **c)** Enrichment analysis for up-regulated genes with a peak on increased accessibility within 1Mb using gene sets from either the “Oncogenic signatures” or “Hallmarks” subsets of the MSigDB database. **d)** Chromatin state of differentially open pan-MM peaks across a developmental range of B-cell types as determined by the Blueprint Epigenomics consortium using ChromHMM (nB: naïve B cells, GCB: germinal center B cell, mB: memory B cell, tPC: tonsil PC and MM: myeloma PC). Enhancers with *de novo* formed peaks in myeloma are indicated. **e)** Radial plot of Homer motif analysis displaying the top 50 over-represented TF motifs in *de novo* myeloma enhancers.

The 2,581 unique differentially accessible regions are predicted to regulate 1,354 unique DEG in at least one genetic subgroup (457 in CCND1, 437 in HD, 914 in MAF and 622 in MMSET). Of note, 977 of the 2,581 (38%) regions were predicted to regulate more than one DEG, while 962 of 1,354 DEG (71%) were predicted to be regulated by more than one differentially accessible region often in more than one genetic subgroup, suggesting a process of chromatin accessibility-based convergence evolution.

Several of the DEG linked to these distal differentially accessible regions have been implicated in myeloma pathogenesis (e.g., *HGF, DKK1, UCHL1*; **Figure 3b and Suppl Fig 2a & b)**. In two myeloma cell lines tested, these same regions are marked by H3K27ac, a histone hallmark of active enhancers, and therefore in terms of chromatin status they can be considered *bona fide* enhancers.

These candidate enhancer-regulated DEG showed overall enrichment for previously defined myeloma transcriptional signatures **(Suppl Fig 3a)** thus further supporting the role of the myeloma ‘enhanceome’ in regulating the oncogenic transcriptome. They were also notable for enrichment of the oncogenic Ras pathway, activated in 40% of MM patients (Chapman et al., 2011; Lohr et al., 2014; Walker et al., 2015) and activation of the Hedgehog pathway, previously implicated in the regulation of *CCND1* and *CCND2* (Katoh and Katoh, 2008) **(Fig 3c)**. Compared to upregulated genes not associated with regions of increased accessibility, there was an over-representation of genes marked by H3K27me3 and genes regulated by members of the Polycomb repressive complex in several cell types **(Suppl Fig 3b)** consistent with activation in myeloma PC of Polycomb repressed transcriptional programmes. We also noted an enrichment for genes involved in ectodermal cell types, such as skin and neuronal cells types, particularly in the HD and MMSET subgroups, suggesting a potential, previously unrecognised role of such genes in myelomagenesis **(Suppl Fig 3b & c)**.

### Developmentally ‘re-commissioned’ and *de novo* formed enhancers in MM

Next, we sought to gain insights into the developmental origins of the MM over-accessible candidate enhancers that are predicted to regulate differentially expressed genes in our cohort of 30 myeloma PC. For this purpose, we tracked the candidate enhancer chromatin status, as defined by combinatorial enrichment of histone marks (ChromHMM states(Hoffman et al., 2013)), across different mature B lineage cells i.e., naïve, germinal center (GCB), memory B cells and tonsil PC and myeloma PC, as previously generated by the Blueprint Genomics consortium (Stunnenberg et al., 2016) **(Figure 3d)**.

Considering that an active enhancer requires combined enrichment of H3K27ac with H3K4me1/3 (Dorighi et al., 2017), we identified 254 regions with regulatory potential over 201 genes differentially expressed in our cohort of 30 myeloma PC that are uniquely and only present in myeloma PC **(Figure 3d)**, i.e., they are *de novo* formed (as is the case for example for the enhancers predicted to regulate *HGF* and *UCHL1* shown above). Transcription factor motif enrichment analysis of these 254 regions identified IRF and MEF families amongst others as possible leading transcriptional regulators of their activity **(Figure 3e)**. Indeed, we identified, a smaller number of regions that are active in one or more B lineage cells but inactive in ND PC are activated, i.e., ‘re-commissioned’ in myeloma PC.

Finally, a higher proportion of putative myeloma enhancers were within 1Kb of Polycomb repressed chromatin in GCB cells and tonsil PC than naïve or memory B-cells (14.2% and 14.0% vs 7.7% and 6.0% respectively, **Suppl Fig 3d**) suggesting that myeloma enhancers in part reactivate a transcriptional programme repressed in GCB and PC **(Figure 3d).**

Thus, biochemical annotation of distal over-accessible profile identifies enhancers that are myeloma PC unique or developmentally inherited.

### Transcription factor ‘rewiring’ is associated with novel myeloma dependencies and prognostic variables

Next, we employed ATAC-seq footprinting on all accessible chromatin to identify predicted association of DNA binding factors (Buenrostro et al., 2013) with chromatin across myeloma and ND PC. In total, 116 of 253 expressed TF **(Figure 4a)** displayed higher or similar predicted binding frequency in myeloma subgroups compared to normal PC, and included TF such as XBP1, IRF4 and PRDM1 known to regulate myeloma transcriptomes, but also TF such as MEF2C and ZNF384 which have not been previously linked to myeloma biology (**Figure 4b & c).** Another 115 TFs were predicted to bind to chromatin in at least one MM subgroup but not in ND PC, including established, subgroup-specific oncogenic drivers (e.g. MAF). Two-thirty two of these 115 TF were predicted to be exclusively active in individual subgroups, with ISL2, a neural TF(Thaler et al., 2004) not previously linked to MM, showing activity solely in the HD subgroup **(Figure 4a-c**).

**Figure 4.**
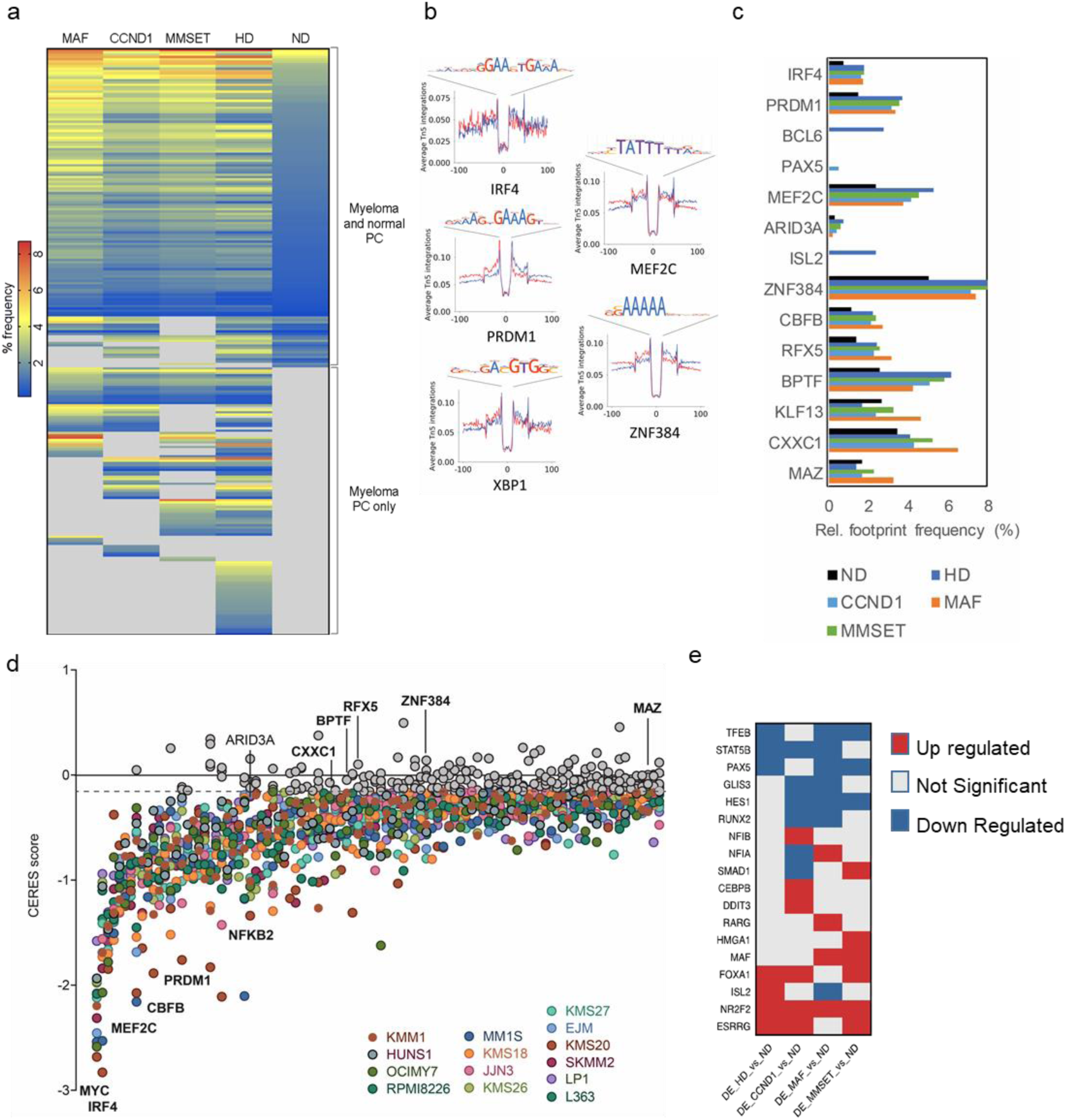
‘Rewiring’ of transcription factors underpins aberrant regulatory gene networks in MM. **a)** Heatmap representation of relative frequency of TF footprints across different MM genetic subgroups. TF are clustered based on their presence in both normal donor and myeloma PC (top), or different myeloma subgroups (bottom). **b)** Footprints of indicated TFs identified in HD myeloma PC as determined by ATAC-seq. **c)** Difference in relative frequency of active TFs between different myeloma subgroups and normal donor PC. **d)** TF dependency analysis by DepMap (14 myeloma cell lines representing CCND1, MAF and MMSET genetic subgroups; color-coded). Of the 248 TF shown, 181 are predicted to generate dependency, the top 100 of which are shown here. Examples of established and novel (bold) dependencies are indicated. In this analysis, dependency is defined as CERES score < −0.1 (horizontal dotted line) in at least 3/14 myeloma cell lines. **e)** TF that are differentially expressed between ND PC and myeloma genetic subgroup PC.

CRISPR/Cas9 depletion of 248/253 TF as retrieved from the DepMap database suggested myeloma cell dependency on 73% TF in at least 3/14 myeloma cell lines **(Figure 4d)**, and confirmed MM cell dependency on the TF *MEF2C* and *ZNF384* identified by ATAC-seq footprinting **(Figure 4c & d and Suppl Fig 4a).** Interestingly only 10% (18/181) of these TF are differentially expressed in one or more myeloma subgroups compared to normal PC **(Figure 4e)**. This is consistent with a pattern of TF activity ‘re-wiring’ in myeloma PC that does not necessarily require transcriptional deregulation of the TF themselves.

For each MM subgroup and ND PC we built TF regulatory gene networks based on weighted frequency of binding and level of expression **(Figure 5a).** In general, compared to ND PC, we observed a higher number of active TF in all myeloma subgroups. These formed higher density regulatory connections with other TF and displayed auto-regulatory loops **(Suppl Fig 4b)**, commensurate with increased binding frequency for >90% of TF in each myeloma subgroup **(Suppl Figure 4a).** Focusing on the MAF-translocated subgroup, using ChIP-seq we obtained the cistrome of oncogenic MAF in the MAF-translocated myeloma cell line MM.1S **(Suppl Fig 4c).** MAF binding was enriched at TSS/promoters and intergenic areas **(Suppl Fig 4d)** and motif analysis identified highly significant enrichment for the MAF motif **(Suppl Fig 4e)** and motifs of IRF1-4, NRF2/NFE2L2, ATF3, BACH1 **(Suppl Fig 4f**); these TF are also predicted to be active within the MAF subgroup regulatory gene network.

**Figure 5.**
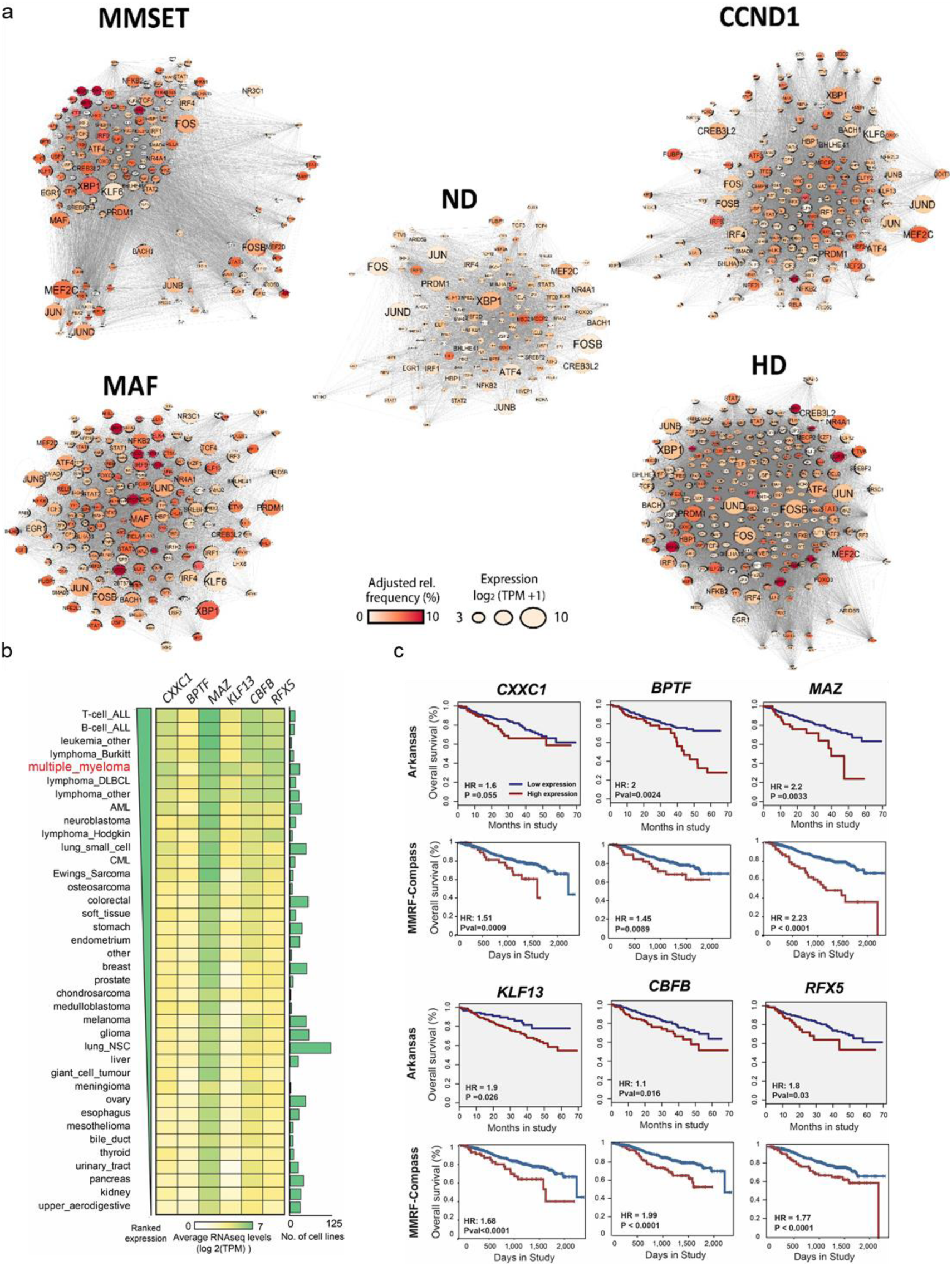
TF regulatory gene networks and novel biological and clinical insights. **a)** TF regulatory gene networks per myeloma subgroup and normal donor PC, as inferred from footprinting analysis. TF are weighed by relative TF binding frequency (color) and expression (node size). **b)** Heatmap of ranked expression of *MAZ, CXXC1, KLF13, BPTF, RFX5* and *CBFB* across >1000 human cancer cell lines (CCLE dataset). Multiple myeloma cell lines are highlighted in red. Bar plot depicts the number of cell lines per cancer group. **c)** MM patient stratification based on *MAZ, CXXC1, KLF13, BPTF, RFX5* and *CBFB* expression (red, high; blue, low) and analysis of overall survival using the Multiple Myeloma Arkansas (n=414) and the MMRF Compass Dataset (n=745). HR: hazard ratio.

Six TF, CXXC1, BPTF, MAZ, KLF13, CBFB and RFX5 were identified as highly connected in all myeloma subgroups but also ND PC, thus exemplifying the process of existing TF ‘re-wiring’ in MM. These TF, none of which have been previously linked to MM, share another three notable features: they demonstrate dependency on DepMap **(Figure 4d)**; display highest expression in multiple myeloma cell lines compared to >1000 other cancer cell lines; and their over-expression has a pronounced impact on survival in two independent myeloma patient cohort datasets **(Figure 5b & c)**. Together, this multi-layered approach reveals novel TF dependencies and prognostic variables in MM.

### Discovery and characterisation of the *CCND2* super-enhancer

Next, by exploiting insights gained from the combined chromatin accessibility-transcriptome profiling, we sought to identify and characterise the regulatory mechanisms of *CCND2* overexpression in MM.

LF5 of MOFA completely separated *MAF*-translocated from *CCND1*-translocated samples **(Figure 6a & b)**, placing extreme opposite weights on the expression of *CCND2* and *CCND1* respectively. This factor places high importance on a set of open-chromatin regions upstream of *CCND2*, linking them to enhanced expression of *CCND2* and transcripts upstream but not downstream of *CCND2* **(Figure 6c)**. These regions were also accessible in the *CCND2*-expressing MMSET and HD samples but minimally accessible in ND PC or in *CCND1*-translocated cases, which lack expression of *CCND2* **(Figure 6d & e).** Further, H3K27ac histone mark enrichment in *MAF*-translocated MM.1S myeloma cells identified the region of interest as a *bona fide* super-enhancer **(Figure 6d).**

**Figure 6.**
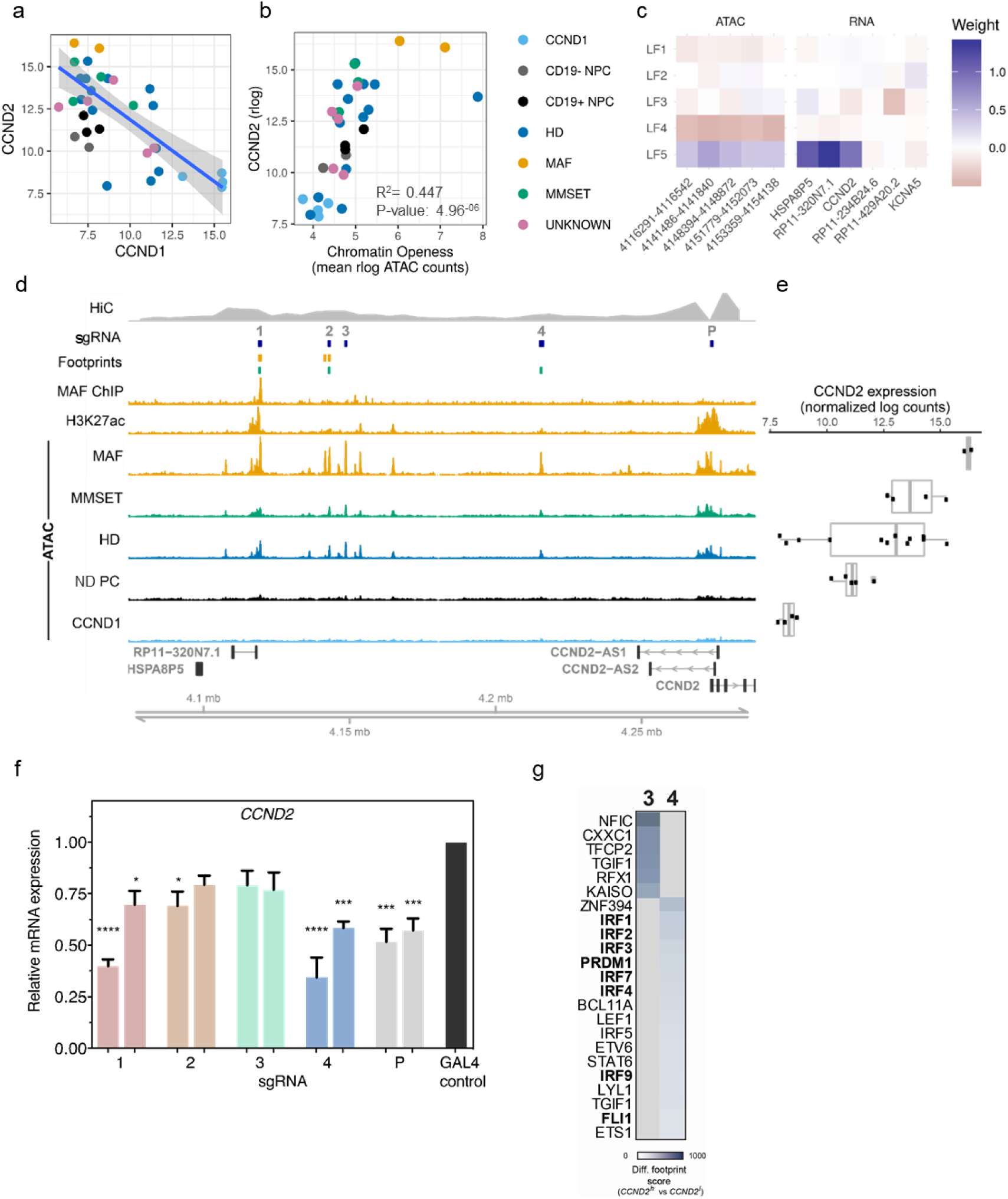
A novel super-enhancer regulates transcription of *CCND2* in myeloma. **a)** Inverse correlation between expression of *CCND1* and *CCND2* across all MM and normal PC samples. Best linear regression fit is indicated as blue line with standard error in grey. **b)** Average chromatin accessibility of regions upstream of *CCND2* and *CCND2* gene expression across all samples in this study. **c)** Loadings from MOFA analysis for 5 regions and 6 genes within 1Mb of *CCND2* that were selected for MOFA analysis for latent factors 1-5. Both regions and genes ordered in a 5’->3’ direction. **d)** Genomic tracks visualization of the *CCND2* region. From top to bottom: Hi-C signal for interactions with *CCND2* promoter in GM12878 B cells; target location of sgRNAs designed for the CRISPRi experiment; Predicted footprints for MAF TF in MAF-translocated (orange) and MMSET-translocated (green) samples; ChIP-seq signal against MAF and H3K27ac in MM1.S cells; normalised ATAC-signal in each subtype, ordered by mean *CCND2* expression. **e)** *CCND2* expression as assessed by RNA-seq in samples across different MM subgroups and normal donor PC. **f)** *CCND2* expression as assessed by qPCR 4 days after CRISPRi of *CCND2* super-enhancer and promoter regions. Two sgRNAs were employed to target the promoter (P) and each of four major peaks (1-4) of the *CCND2* super-enhancer, as shown in (d), and compared with a non-targeting control (Gal4). n=3, *p<0.05, *** p<0.001, **** p<0.0001, mean and standard error of the mean are shown. **g)** Heatmap illustration of TF motifs significantly enriched in *CCDN2*^high^ versus *CCND2*^low^ HD samples, as identified by differential footprinting analysis. Differential footprints were identified in peaks 3 & 4 (see Fig 6d).

Consistent with it being a ‘re-commissioned’ enhancer, the region of interest is Polycomb repressed in GCB and PC but active in naïve and memory B cells **(Suppl Fig 5a)**, consistent with *CCND2* being expressed in naïve and memory B cells but not GCB cells or PC **(Suppl Fig 5b).**

Using Hi-C genome data generated in EBV transformed B cells (Rao et al., 2014) which resemble PC in that they can secrete Ig, we identified exclusive, high frequency long range interactions of the *CCND2* promoter with the upstream accessible clusters of the putative enhancer **(Figure 6d and Suppl Fig 5c)** further validating the region of interest as a *bona fide CCND2* enhancer.

In a complementary approach, we employed KRAB-dCAS9 CRISPRi in MAF-translocated cells to repress activity of four prominent constituent peaks (1-4; **Figure 6d**) which engage in high frequency interactions with the *CCND2* promoter. As expected, targeting of a promoter accessibility peak resulted in a significant decrease of *CCND2* expression, while in the *CCND2* enhancer, the most pronounced effect, similar to that of the promoter peak, was conferred by targeting of the proximal peak 4 and distal peak 1 accessibility areas **(Figure 6f).** Notably, peaks 1 and 4 but not accessibility peaks in between, are Polycomb-repressed in GCB and PC but active in naïve and memory B cells **(Suppl Fig 5a).** Thus, as well as by experimental means (i.e., CRISPRi), the relative importance of peaks 1 and 4 is also validated from a developmental perspective.

Having dissected the *in cis* regulatory mechanisms of *CCND2* transcriptional regulation we proceeded with the characterisation of *trans* factors involved in this process. Previous work showed that MAF binds to *CCND2* promoter *in vitro*(Hurt et al., 2004) thus providing some insight into how *CCND2* is regulated in the MAF genetic subgroup. Importantly, our ChIP-seq analysis in MM.1S myeloma cells shows that MAF binds to the *CCND2* enhancer *in vivo* **(Figure 6d)**, thus consolidating the role of MAF as a critical regulator of *CCND2* over-expression in *MAF*-translocated MM cells. This finding provides also insights into *CCND2* regulation in the MMSET genetic subgroup which is characterised by lower level expression of both *CCND2* and *MAF* than the MAF subgroup (Hurt et al., 2004) **(Figure 6e and Suppl Fig 5d).** Since chromatin accessibility signal in the MMSET subgroup is also lower than in the MAF subgroup **(Figure 6d)**, together, these findings are consistent with the notion that *CCND2* enhancer transcriptional activity in the MAF and MMSET subgroups is MAF dosage-dependent.

To identify TF potentially regulating *CCND2* enhancer activity in *CCND2*-expressing HD MM which lack expression of MAF, we performed differential footprinting analysis in CCND2^hi^ vs CCND2^lo^ HD samples **(Figure 6g and Suppl Fig 5e & f).** In addition to TF known to be implicated in MM (IRF4, FLI1)(Jin et al., 2018; Shaffer et al., 2008), we also identified a potential regulatory role for TF previously not linked to MM (e.g., CXXC1, ZNF394, and IRF3).

Finally, we explored activity of the *CCND2* super-enhancer in chronic lymphocytic leukemia (CLL), the most common B cell lineage malignancy. High CCND2 expression has been previously documented in CLL malignant B cells residing in proliferation centres(Igawa et al., 2011), the anatomical areas in secondary lymphoid organs where malignant B cells receive survival and proliferative signals(Zhang and Kipps, 2014). CLL malignant B cells express significantly higher *CCND2* than *CCND1* **(Suppl Fig 6b)** and as in *CCND2*^hi^ myeloma PC, we found that the same *CCND2* super-enhancer is active in CLL B cells **(Suppl Fig 6a)** thus expanding the importance of the *CCND2* enhancer in a wider range of B cell lineage malignancies.

## Discussion

Here, by generating and integrating complementary datatypes we dissect how chromatin dynamics link MIE with their oncogenic transcriptomes. In addition, we identify the candidate ‘enhanceome’ for each of the main myeloma genetic subgroups and by charting their corresponding TF regulatory networks we provide novel functional and clinical insights.

Enhanced overall accessibility in all myeloma genetic subgroups is our first fundamental observation underpinning all subsequent findings. It was made possible by inclusion in the study of the myeloma PC normal counterparts, i.e., normal bone marrow PC. This contrasts with a pan-cancer chromatin accessibility profiling which was defined without reference to normal tissue of origin(Corces et al., 2018) and it also differs from previous work in which chromatin accessibility in MM was defined with reference to in vitro generated PC(Jin et al., 2018). These, functionally and from the regulatory genome standpoint would be expected to significantly diverge from in vivo generated bone marrow PC.

Having obtained MM samples from all main molecular genetic subgroups and a faithful reference population, we used an unsupervised, machine-learning approach to demonstrate the power of combined chromatin accessibility and transcriptome analysis to categorise MM versus ND PC according to MIE. This was possible for all IgH-translocated subgroups, but not for HD MM, likely because the latter represents an inherently biologically heterogenous group. Whether MIE themselves are directly involved in or impose chromatin accessibility changes is an open question. It the case of MAF-translocated MM for instance, it could be envisaged that ectopically expressed MAF acts as a pioneer TF (Mayran and Drouin, 2018) with ability to initiate chromatin de-condensation. The finding that MAF translocated samples demonstrated the most pronounced divergence from ND PC in MOFA and had the largest number of distal, differentially accessible chromatin regions would be consistent with such a scenario.

Moreover, our findings show that the regulatory potential of over-accessible chromatin regions is unequally distributed between proximal and distal regulatory elements; the latter are more likely to regulate the myelomagenic transcriptome, highlighting the primacy of these putative enhancers in shaping myeloma cell biology. Importantly, we also show that these distal elements correlate with genes differentially expressed between myeloma and normal PC within the boundaries TADs thus satisfying the requirement that enhancer-candidate gene target interactions are delimited in the 3D genome space (Robson et al., 2019). Another level of functional validation of these candidate enhancers is provided by the finding that genes differentially expressed in MM and predicted to be regulated by candidate enhancers are highly enriched in previously defined myeloma transcriptome signatures. Importantly, putative enhancers are also predicted to regulate genes downstream of oncogenic RAS pathway which is activated by secondary gain-of-function N- or K-RAS somatic mutations in 40-50% of MM cases(Chapman et al., 2011; Lohr et al., 2014), thus highlighting the ability of our approach to also reveal chromatin traces of secondary driver genetic events. At individual gene level, we identified myeloma PC-specific, putative enhancers marked by H3K27ac for *DKK1, HGF* and *UCHL1*, linked to their aberrant over-expression in MM. *DKK1* is a soluble Wnt pathway antagonist that inhibits differentiation of osteoblasts when secreted by myeloma PC thus contributing to development of myeloma bone disease(Qiang et al., 2008); secreted *HGF* in a autocrine and paracrine manner promotes myeloma cell proliferation and migration as well as angiogenesis(Gambella et al., 2015); while, the de-ubiquitinase *UCHL1* is required for myeloma dissemination and high expression levels at diagnosis predict early disease progression(Hussain et al., 2015). Another notable feature mostly restricted to HD MM is the activation, via putative enhancers, of genes involved in neurogenesis. The functional significance of this novel finding is not clear and requires further investigation. We note that recent work demonstrated ectopic activation of neural genes in different cancers that engage in functional cross-talk with the central and peripheral nervous system(Magnon et al., 2013; Zhao et al., 2014).

By taking advantage of biochemically defined enhancer functional states in the whole spectrum of B lineage cells we validated over-accessibility regions corresponding to 830 candidate enhancers in myeloma PC as functionally active enhancers marked by H3K27ac. The candidate enhancers of *HGF* and *UCHL1* exemplify *de novo* formed enhancers, i.e., not present at any stage of late B lineage development, while the *CCND2* enhancer provides an example of ‘re-commissioned’ enhancer, i.e., active in myeloma but not in normal PC (see below). In many cases, recommissioning entails activation in myeloma PC of Polycomb-imposed poised transcriptional states in GCB cells and normal PC.

As well as discovery of critical *cis* regulatory elements, ATAC-seq also affords the opportunity for inferring TF binding to chromatin through footprinting and motif analysis. Through this process we identified the compendium of TF that are predicted to regulate myeloma transcriptomes. These include TF such as *IRF4* and *PRDM1* that define PC identity and were previously shown to sustain proliferation and survival of malignant PC. However, a more notable finding is that the majority of *trans* factors that display increased binding frequency to chromatin are already active in ND PC and their expression is not deregulated in MM. Such TF engage at higher frequency interactions with other TF than in ND PC and are more likely to self-regulate, properties that have been associated with enhanced regulatory potential (Saint-Andre et al., 2016). These properties, as revealed through construction of TF regulatory gene networks, led to novel insights with functional and clinical implications. For example, we identified TF such as MEF2C and ZNF384 to which myeloma cells appear to be addicted. In addition, the case of CXXC1, BPTF, MAZ, KLF13, CBFB and RFX5, six TF with unknown function in MM and incompletely studied in general, also highlights the strength of our functional epigenomics approach. None of these TF is differentially expressed in MM, yet they display increased predicted regulatory potential across all myeloma subgroups, likely reflecting their high expression levels in myeloma cells compared to other cancers. Moreover, all six TF demonstrated prominent myeloma cell dependency and were found to strongly predict prognosis. Future research will further validate and define the role of these and other novel TF identified herein in myeloma biology.

Extensive genetic heterogeneity and diversification in MM poses significant therapeutic challenges. One of the striking features of myeloma biology is the early observation that a dichotomous over-expression of the cell cycle regulators *CCND1* and *CCND2* over-arches genetic diversification (Bergsagel et al., 2005). While in the majority of MM cases overexpression of *CCND1* can be explained by somatic structural variants i.e., juxtaposition to IGH@ enhancer or chr11q25 gain(Shah et al., 2018), no such aberrancies account for overexpression of *CCND2*. Our identification and functional validation of the distal *cis* and *trans* regulators of *CCND2* expression addresses this gap in the biology of MM.

Using orthogonal approaches (genetics, chromatin accessibility and ChromHMM maps, 3D genome analysis, epigenetic silencing, transcriptome profiling and correlation with *in vivo* developmental trajectory) we clearly demonstrate the enhancer-gene target relationship between *CCND2* and its proposed enhancer.

This allowed further insights into how activity of this enhancer is regulated in different myeloma genetic subgroups. While expression of *MAF* is highest in the MAF subgroup as a result of its juxtaposition to the powerful IgH@ enhancer, in MMSET MM expression of *MAF* is lower and previously shown to be regulated by Fos in response to activated mitogen-activated protein kinase pathway Annunziata et al., 2011). It is likely that these differences in MAF dosage account for the strongest chromatin accessibility signal at the MAF-bound *CCND2* enhancer and higher level of *CCND2* expression in MAF-translocated MM. Finally, we identified known and novel candidate *in cis* regulators of the *CCND2* enhancer in *CCND2*-expressing HD MM which will require additional experimental validation.

The *CCND2* enhancer is also a prime example of ‘recommissioned’ enhancer i.e., repressed by Polycomb in GCB and PC but active in naïve and memory B cells. Activation in myeloma PC of Polycomb repressed distal regulatory elements and of genes regulated by them permeates the myeloma ‘enhanceome’ and likely is an important mechanism for activation of transcriptional programmes repressed in normal PC. Interestingly, the same enhancer with the same developmental epigenomic features regulates expression of *CCND2* in CLL cells, a finding that tallies with the notion that CLL originates either from a naïve or memory rather GCB cell(Beekman et al., 2018).

Precise identification and functional dissection of the *CCND2* super-enhancer and indeed of all enhancers that regulate genes critical for cancer biology is not only important for the delineation of molecular mechanisms that drive oncogenesis. It is also a pre-requisite for enhancer-focused, epigenome-directed therapeutic interventions. As previously shown for the super-enhancers regulating Hb globin genes, this can be facilitated by technologies that allow unbiased, de novo analysis of DNA binding factors and chromatin modifiers that are enriched in the enhancer of interest (Liu et al., 2017).

In sum, with reference to cell of origin, we linked primary myeloma genetic drivers to their oncogenic transcriptomes through a multilayer approach centered on chromatin dynamics. This allowed identification and integration into regulatory networks of *cis* and *trans* regulatory chromatin elements, facilitated gain of novel pathogenetic insights with biological and clinical implications and provides a resource for further biological exploration in MM. Our discovery and functional dissection of the critical for the biology of MM *CCND2* enhancer is a prime example of the power of this approach and affords opportunity for enhancer-based therapeutic approaches.

**Table 1.** Antibodies.

| Antibody | Purpose | Fluorophore Channel |
| --- | --- | --- |
| CD138 | positive | FITC |
| CD319 | positive | PE |
| CD27 | positive | PECy7 |
| CD2<br>CD3<br>CD14<br>CD16<br>GPA | negative | Alexa647 |
| CD19 | Pos & Neg | APC efluoro780 |
| CD45 | Positive | Alexa700 |
| CD38 | positive | Alexa405 |
| 7AAD | negative | 7AAD |
| CD56 | FACs MM | PECy7 |
| CD38 | FACs MM | BV421 |

**Table 2.**
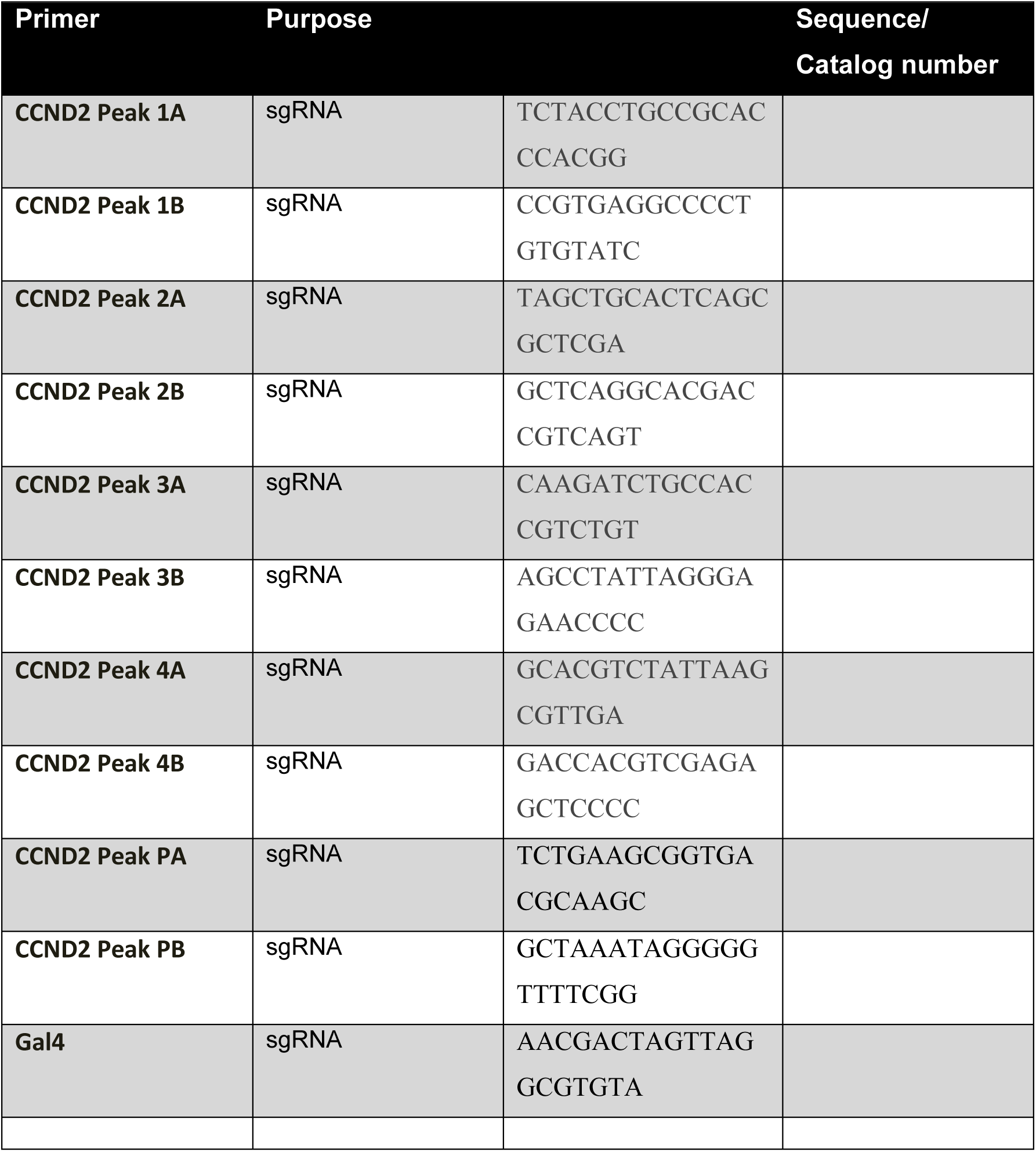
Primers.

## Acknowledgments

NT, VC, XX and KP were supported by Bloodwise, AlK was supported by Kay Kendall Leukaemia Fund and Imperial NIHR Biomedical Research Centre. We acknowledge support from Imperial NIHR Biomedical Research Centre, Cancer Research UK Experimental Cancer Medicine Centre, LMS/NIHR Flow-cytometry Facility, Imperial NIHR Biomedical Research Centre Genomics Facility.

## Materials and Methods

### Patient samples

Bone marrow (BM) samples were obtained after written informed consent and research ethics committee approval (Research Ethics Committee reference: 11/H0308/9).

Patient BM aspirates were subjected to red cell lysis. Multiple myeloma plasma cells were purified by two rounds of CD138 immunomagnetic selection (Miltenyi Biotech) following the manufacturer’s instructions. Pre and post selection purity was assessed by FACs analysis (BD LSR-Fortessa) using CD138, CD45, CD19, CD56 and CD38 panel antibodies (Table 1). Purified cells were immediately processed for ATAC-seq and RNA-seq.

### Normal donor samples

BM-MNCs from BM aspirates were isolated by ficoll (Histopaque, Sigma). The MNCs were pre-cleared of T cells and Monocytes by consecutive immunomagnetic negative selection (CD3 and CD14-EasySep StemCell Technologies) following manufacturer’s instructions. The sample was stained and sorted for positive CD138, CD319, CD27, CD45 and CD38, and negative for CD2, CD3, CD14, CD16, GPA and 7AAD. (FACSAriaII, BD Biosciences) and CD19^+^ or CD19^-^. The sorted cells were immediately processed for ATAC-seq and RNA-seq.

### Fluorescence in situ hybridisation (FISH)

FISH was undertaken using a panel of 4-7 probe sets targeting regions of common cytogenetic abnormalities in multiple myeloma (Kreatech Diagnostics, Amsterdam, The Netherlands). Interphase cells were dropped onto a glass slide and dried briefly before fixation in situ. Hybridisation was performed according to manufacturer’s protocols. The panel consists of two probes (13q14 and 13qter) to detect deletion and monosomy of chromosome 13, a locus specific probe to detect deletion of TP53 (17p13), two probes on chromosome 9 and 15 to detect hyperdiploidy and a dual colour, break-apart probe to detect rearrangements of IGH (14q32). Rearrangements of IGH were further investigated with IGH/CCND1, IGH/MAF and IGH/FGFR3 dual colour, dual fusion probes. The upper threshold for normal results are according to probe type (dual colour break apart 5%; quantitative 5%; dual colour dual fusion 2%). In all cases, a minimum of 50 interphase cells were scored by two independent analysts.

### Cell lines

The human multiple myeloma cell line (MMCLs) MM.1S (ATCC, Manassas, VA, USA) was cultured in RPMI1640 media (Sigma, UK) and 20% FBS (Life Technologies). JJN3 cells(DSMZ, Germany) were cultured in 40/40% DMEM/IMEM medium (Sigma, UK), and 20% FBS.

HEK 293T cells were cultured in DMEM (Sigma, UK), 10% FBS (Gibco). All cell lines were maintained at 37°C and 5% CO2 and the growing media was supplemented with 1% penicillin/streptomycin (Sigma, UK) and 1% L-glutamine (Sigma, UK).

Testing for mycoplasma presence was performed every 4 weeks.

### ATAC-seq

ATAC-seq was performed as described(Buenrostro et al., 2013). Briefly, 50,000 purified plasma cells, myeloma plasma cells or cell lines, were washed with cold PBS (Sigma, UK) at 500g at 4^°C^ for 5 min. The cells were resuspended in 50 μL of cold Lysis Buffer (10 mM Tris-HCl, pH 7.4, 10 mM NaCl, 3 mM MgCl_2_, 0.1% IGEPAL CA-630) and washed at 500g at 4^°C^ for 10 min. The nuclei were subjected to transposase reaction for 30 min at 37^°C^; termination of the reaction and DNA purification was performed using a MiniElute Kit (Qiagen) and eluted twice with 10 μL. The purified DNA was amplified as described before with NEBNext High-Fidelity 2x PCR Master Mix (New England Biolabs). The PCR amplified product was cleaned twice with (0.9X) AMPure beads (Beckman). The quality of the libraries was assessed with the Bioanalyzer High Sensitivity DNA kit (Agilent). The libraries were quantified using the NEBNext Library Quant Kit for Illumina (New England Biolabs) on a StepOne Plus Real-Time PCR (Applied Biosystems). The libraries were sequenced at the Genomics Facility at ICL using the Illumina HiSeq 4000 platform to obtain paired-end 75bp reads.

### RNA-extraction and cDNA synthesis qPCR

Purified plasma cells, myeloma plasma cells or cell lines (100,000), were washed with cold PBS (Sigma, UK) at 500g at 4^°C^ for 5 min. Total RNA was isolated using the Nucleospin RNA kit (Macherey-Nagel) and quantified by Nanodrop Lite (Thermoscientific). cDNA was synthesized with RevertAid cDNA synthesis kit (Thermoscientific). qRT-PCR for was performed with Taqman probes (Applied Biosystems) using StepOne Plus Real-Time PCR (Applied Biosystems). Gene expression was normalized to the expression of *GAPDH* (Table 2).

### RNA-seq

Total RNA was isolated using the Nucleospin RNA kit (Macherey-Nagel) and quantified using the Qubit RNA Assay kit (Life Technologies) and RNA quality was assessed on the Bioanalyser using the RNA pico kit (Agilent). Total RNA libraries were prepared by removing the ribosomal RNA with NEBNext rRNA depletion kit (New England Biolabs) and NEBNext Ultra II Directional RNA Library Prep kit for Illumina (New England Biolabs), following manufacturer’s instructions. Library quantity was determined using the Qubit High Sensitivity DNA kit (Life Technologies) and library size was determined using the Bioanalyser High Sensitivity DNA kit (Agilent). Libraries were diluted to 2nM and sequenced using the Illumina HiSeq 4000 platform the Genomics Facility at ICL to obtain paired-end 75bp reads.

### dCas9-KRAB CRISPRi

Two single guide RNAs (sgRNAs) targeting each peak were designed using the online tool http://crispr.org (Table 2). Two sgRNAs were designed for the *CCND2* promoter as a positive control and the non-targeting sgRNA for Gal4 of the yeast *S.cerevisiae* as a negative control.

The vector used was the inducible Lenti-CRISPR-dCas9-KRABv2. The original LentiCRISPRv2 vector (Addgene 52961, USA) was modified and kindly supplied by Dr Feldhahn’s Lab, Imperial College London. The plasmid was digested with Esp31 (NEB, UK), removing a 2kb stuffer.

The sgRNA oligos were phosphorylated and annealed using the T4 Ligation Buffer (New England Biolabs) and T4 PNK (New England Biolabs). The sgRNAs were then ligated with the digested plasmid using the T4 DNA ligase and buffer (New England Biolabs). Reactions were carried out following the Zhang Lab General Cloning Protocol (Addgene).

Sanger sequencing was used to confirm the exact sequence of the cloned sgRNA.

### Lentivirus production and cell transduction

The dCas9-KRAB constructs were transfected in 293T cells with the 2nd generation lentiviral packaging and envelope plasmids psPAX2 and pMD2.g (Addgene, USA), using CaCl2 2.5M (Sigma, UK) and 2x HEPES-Buffered Saline, pH=7 (Sigma, USA). After 10 hr, cells were incubated with glycerol (15% v/v) (Honeywell, USA) for 3 min. Cells were washed with PBS (Sigma, UK) and then incubated with fresh DMEM medium (Sigma, UK) for 36 hours. The virus was harvested, filtered (0.45μm) and concentrated by ultracentrifugation (23000rpm for 1 h and 40 min, at 4^°C^) 48 h and 72 h post transfection using the Thermo Sorvall Ultracentrifuge, MTX (Applied Biosciences) Transduction of JJN3 MMCL was performed in the presence of 8 μg/ml polybrene (Sigma, UK). On day 3, transduced JJN3 cells were selected with puromycin (5μg/ml), and 48 h later viable cells were sorted on MA900 Multi-Application Cell Sorter (Sony Biotechnology) and cultured in fresh DMEM/IMEM medium (Sigma, UK). Cells were grown for 10 days, before induction with doxycycline (Sigma, UK) topped up daily to a final concentration of 1μg/ml. On day 4 GFP+ cells were FACS sorted for RNA extraction.

## COMPUTATIONAL ANALYSIS

Unless stated human genome version hg38 with alternative contigs removed was used for all analyses, and annotations were taken from Ensembl version 85.

### RNA-seq analysis

#### Read cleaning and filtering

Paired-end RNA-seq reads were adapter trimmed by the DNA sequencing facility at Centre for Haematology, Division of Experimental Medicine Faculty of Medicine, Imperial College London were quality controlled with FastQC version 0.11.3.

#### RNA-seq Quantification

Expression estimates for each sample were obtained using Salmon version 0.11.4 (Patro et al., 2017), using the complete Ensembl 85 transcriptome. Salmon was used with fragment GC bias correction, 100 bootstrap samples and using an auxiliary k-mer hash over k-mers of length 31.

#### Correction of batch effects, normalisation and differential expression

RNA-Seq normalisation and differential expression were performed using DESeq2, version 1.18.1 (Love et al., 2014). Transcript read counts were summarised to per-gene counts as the total in all transcripts of that gene (based on the transcriptome table created in Salmon). For MOFA analysis and visualization, read counts were rlog transformed (blind=True) using the rlog function from DESeq2. Samples were adjusted for sequencing batch accounting for PC and MM subgroup effects using the removeBatchEffects function from limma version 3.34.9 (Ritchie et al., 2015).

For differential expression analysis and inTAD analysis, gene counts for CD19^+^ and CD19^-^ samples from the same donor were collapsed using the function collapseReplicates from the Deseq2 package and R version 3.4.1. As normal PC (NPC) sample RS_4.25 is the only sample passing quality control in its batch (pool4), batch effects for this sample can’t be quantified. It was therefore assigned to pool1 batch since this batch contains the most samples and in this way, the batch effect change on pool1 should be minimum. Additionally, only for inTAD analysis, read counts were rlog transformed (blind=True) and samples were adjusted for sequencing batch accounting for PC and MM effects using the removeBatchEffects function from limma version 3.34.9 (Ritchie et al., 2015). To identify genes with significantly different expression between normal and cancer PCs, a Wald test was performed using DESeq2, on all samples, comparing cancer to normal, accounting for sequencing batch. To identify genes where expression significantly varied between subgroups, a Log Ratio Test (LRT) of subgroup MM vs. PC (condition) accounting for batch was performed using DESeq2 on normal samples, and cancer samples where a cytogenetic classification was available.

Results were designated significant if they had an adjusted p-value(p-adj) less than 0.1 and absolute log2FoldChange greater or equal to 1.5, either between cancer and normal (pan-myeloma genes), or between at least one subgroup and normal (subgroup genes) were kept.

### Annotated and unannotated Transcription Start Sites (TSS)

In order to obtain unannotated TSS present in the PC and MM samples (primary and cell lines), RNA-seq reads were mapped using Hisat v0.1.6(Kim et al., 2015). Stringtie version 1.2.3 (Pertea et al., 2015) was used to assemble mapped reads in conjunction with the reference geneset to generate novel transcripts. Single exon transcripts were removed from this list (under the possibility that they may be eRNA transcripts). The TSS of these novel transcripts were taken to be the first 1bp of the transcript. Annotated TSS were obtained by taking the first 1bp of coding and non-coding genes from Ensembl v85.

Annotated and unannotated TSS sites were transformed into promoter regions by extending 2kb upstream of the TSS, and 100bp downstream to cover the TSS site using Bedtools version 2.22.1(Quinlan and Hall, 2010); these are referred to as promoter sites.

### ATAC-seq analysis

#### Read cleaning and filtering

Raw paired-end ATAC-seq reads were quality controlled using FastQC. ATAC-seq adapters were removed, uncalled bases (N’s) on ends of reads were trimmed using Cutadapt version 1.9.1(Martin, 2011). Only read pairs with both single end fragments remaining were kept. A second quality pass was performed using Sickle version 1.33 (https://github.com/najoshi/sickle), trimming was performed with a sliding window average Phred quality threshold of 30 (without five prime trimming). Only pairs with a minimum of 20 bases in each read were retained.

#### Mapping and calling chromatin accessible peaks

As per the ENCODE ATAC-seq pipeline, the remaining paired-end reads were mapped using Bowtie2 version 2.3.0 (Langmead and Salzberg, 2012) in paired-end mode to the human genome reporting up to 4 alignments per read, with maximum fragment length 2000bp. Processing was performed using pipeline_atac_consensus_balanced_peaks (https://doi.org/10.15131/shef.data.12280904) based on the ENCODE pipeline and recommendations for ATAC-seq processing (https://www.encodeproject.org/atac-seq/) to produce peaks in signals and reads in peaks per sample, using the CGAT-core framework.

Briefly, the pipeline first filters correctly mapped reads. Reads removed using Samtools version 1.3.1)(Li et al., 2009) include: unmapped (read pair or one of the reads), reads failing platform, orphan reads (one of the reads in the pair removed), read pairs mapping to different chromosomes (-F 524 -f 2 Samtools flags). Read pairs non-overlapping reads in RF orientation were removed with Samtools and Bedtools and multimapped reads using the assign_multimapper script from https://github.com/kundajelab/bds_pipeline_modules/blob/master/utils/assign_multimappers.py. Duplicates were marked using Picard Markduplicates version 1.135 (https://broadinstitute.github.io/picard/) and duplicates removed using Samtools. Read pairs were converted into two single end tags and Tags in mitochondrial and non-standard chromosomes were removed. Soft clipped bases re-added and the TN5 added base pairs on the 5’ sites of each read were trimmed. (these are referred to as shifted tags).

Two consensus peak sets were created: the pan myeloma set and the subgroup set. For the pan myeloma set, the tags in each sample were down-sampled so that each sample had the same number of tags. Samples were then pooled into two categories – MM and NPC.

Broad and narrow peaks were called on each pool using MACS2 version 2.1.1.20160309 (https://github.com/taoliu/MACS)(Zhang, Liu et al. 2008) with the following options for narrow peaks:

-g hs -q 0.01 --nomodel --shift -100 --extsize 200 -B --SPMR --keep-dup all --call-summits

And for broad peaks:

-g hs -q 0.01 --nomodel --shift -100 --extsize 200 --broad --broad-cutoff 0.01 --keep-dup all

Narrow and broad peaks were filtered to remove areas of low mappability (Hoffman, Ernst et al.2013),downloaded from http://hgdownload.cse.ucsc.edu/goldenPath/hg19/encodeDCC/wgEncodeMapability/wgEncodeDukeMapabilityRegionsExcludable.bed.gz and “lifted over” from hg19 to hg38(http://genome.ucsc.edu/cgi-bin/hgLiftOver).Peaks in ENCODE blacklist regions (Amemiya et al., 2019)were also removed, downloaded from https://www.encodeproject.org/files/ENCFF419RSJ/@@download/ENCFF419RSJ.bed.gz).

The narrow peaks and broad peaks from each pool, and peaks less than 200 base pairs apart were merge using bedtools.

For the subgroup peak set, samples without a cytogenetic classification were excluded, and the remaining samples downsampled to the new minimum. Tags from samples in the same cytogenetic group (MAF, MMSET, CCND1, HD or ND PC) were pooled and peaks called on each pool as described above.

#### Sample assigned fraction

The sample assigned fraction reflects the proportion of the total reads that are mapped to areas of high accessibility compared with background noise. To calculate it, sample shifted tags were transformed extending their five prime end 100bp upstream and then extending 200bp downstream (by the same values as used in MACS). They were filtered to remove tags overlapping with areas of low and high mappability defined previously. Sample peaks were obtained by using only sample ATAC-seq data with the method defined on the previous section.

The number of sample filtered, extended shifted tags overlapping sample peaks were obtained using bedtools and divided by the total sample shifted tags to obtain the assigned fraction.

#### Annotations of the consensus peak regions

To annotate regions, the R library Annotatr version 1.8.0(Cavalcante and Sartor, 2017) was used, with hg38 annotations from the library TxDb.Hsapiens.UCSC.hg38.knownGene (https://bioconductor.org/packages/3.9/data/annotation/html/TxDb.Hsapiens.UCSC.hg38.knownGene.html). Any region can overlap multiple different types of genomic annotations on both strands but for each region, a particular type is only reported once. In addition to the Peak sets described above, we also obtained annotation for a randomize sample of regions using the function randomize_regions from the R library Annotatr version 1.8.0.(Cavalcante and Sartor, 2017). The sample is taken without allowing overlaps and with per-chromosome regions being maintained (non alternative, random, unknown and mitochondrial chromosomes not used).

#### Quantitation, normalization and differentially accessible peaks

Shifted tags from each sample were transformed to obtain only the initial 1bp and filtered to remove tags overlapping with areas of low and high mappability defined previously. The number tags overlapping each peak in the peak set is reported using Bedtools intersect.

Samples were assigned to batches based on the sequencing run from which they derived. Where a sample was sequenced in more than one run, its merged reads were assigned a batch that indicated both runs. All samples from a batch that contained only a single sample passing quality control were pooled into a single batch and used as the reference batch for correcting the others.

For MOFA analysis, inTAD analysis and visualization, read counts in the pan myeloma set were rlog transformed and normalized using the rlog function from DESeq, and batch effects removed accounting for PC and MM subgroup effect (MOFA and visualization) or PC and MM effect (inTAD analysis) using the removeBatchEffect function from limma.

To identify differentially accessible peaks between normal and cancer samples, read counts for the pan myeloma peaks were used. Read counts for CD19^+^ and CD19^-^ samples from the same NPC donors were collapsed using collapseReplicates. Significance was determined by Wald test using DESeq2 and the design formula ∼ batch + condition. Outlier removal using Cook’s distance was disabled. Regions were designated as significantly changed if they had a Benjamini-Hochberg corrected p-value of < 0.1 and a log_2_ fold change of at least 1 between cancer and normal.

To identify peaks differentially accessible between subtypes, reads counts for the subtype consensus peak set from samples with a cytogenetic annotation were used (22 Myeloma samples, 5 normal, before collapsing). Read counts for CD19^+^ and CD19^-^ samples from the same NPC donors were collapsed using collapseReplicates. To determine regions that differed between subtypes, a differential analysis was performed using a Log Ratio Test accounting for batch with the design ∼ batch + subtype. Outlier detection using Cook’s distance was used with default thresholds. Significant regions were those having adjusted p-value (p-adj) less than 0.1 and absolute log_2_FoldChange >1 for any MM subgroup vs. NPC comparison.

### Characterising candidate enhancers and regulated genes

To obtain candidate enhancers, we took the set of differentially accessible regions that were more accessible either in cancer (pan-MM enhancers) or in at least one subtype (subtype enhancers) than in NPC. We removed any regions that overlaped the unannotated and annotated promoter site using Bedtools.

We associated these more accessible regions with genes within 1Mb that had significantly differential expression either in cancer or in the appropriate subtype.

### Enrichment analysis

Enrichment analysis was carried out using gene sets from either MSigDB or Enrichr. MSigDB gene category membership was obtained from the msigdbr package (Liberzon et al., 2015). Enrichr gene-category relations were obtained using the Enrichr API(Kuleshov et al., 2016). Gene set enrichment was calculated using the goseq R package version 1.36.0(Young et al., 2010), correcting for gene length. p-values for over-representation were corrected using the Benjimini-Hochberg procedure. For enrichment of differentially expressed genes associated with differentially accessible regions, either all genes not filtered by DESeq, or only differentially expressed genes were used as a background as noted in the text.

### Correlation of gene-expression with peak accessibility in the same TAD

In order to measure the correlation of gene expression with peak accessibility in the same TAD, InTAD version 1.2.3 was used (Okonechnikov et al., 2019). As input, the rlog transformed read count values of differentially expressed genes and differentially accessible non-TSS peaks in the pan-myeloma analysis within a maximum distance allowed between the peaks and genes was 1Mb. Additionally, interactions were restricted to TAD locations from the cell line GM12878 (Rao et al., 2014). Peaks and genes were correlated using Pearson correlation coefficient and p-values were corrected using the findCorrelation method from the inTAD package (adj.pval parameter set to TRUE).

### Modelling effect of differential promoters and candidate enhancers on gene expression

Differentially regulated promoters were obtained by intersecting the TSS set described above with differential peaks from the pan-myeloma peak sets using bedtools intersect. The number of peaks of increased accessibility within 500kb of differential gene TSSs was calculated by subsetting the differential pan-myeloma peaks after removal of TSSs to only those more open in myeloma, and then counting overlaps with the TSS regions around each gene extended by 500,000nt in each direction, again using bedtools intersect. Logistic regression was performed to predict identity of significantly upregulated using presence of promoter of overlapping peak of increased accessibility and the count of other peaks of increase accessibility within 500kb using R function glm, with a binomial family and logit link function.

### Multi-Omics Factor Analysis (MOFA)

#### MOFA Inputs

For ATAC-seq, the rLog transformed, normalized and batch corrected tag count for the pan-myeloma peak set were obtained for all samples (described above). Regions corresponding to sex chromosomes (chrX and chrY) were and regions overlapping multi-exon TSS (see above) were removed. This yielded 273,216 remaining regions. The overall variance per peak was calculated and only the 5000 peaks with the highest variance were selected.

For RNA-seq, the rLog transformed, normalized and batch corrected RNA-seq read counts for all samples were obtained (described above). Genes on sex chromosomes were removed from the table. The variance per gene was calculated and only the 5000 genes with highest variance.

#### Training

Using the R library MOFAtools version 0.99.0(Argelaguet et al., 2018), the ATAC-seq and RNA-seq tables were input to MOFA. The default data and model options were used and “Gaussian” data was selected both for ATAC-seq and RNA-seq. The training options used that were different to the defaults were: dropping factor threshold of 0.01, 10,000 maximum iterations, minimum tolerance convergence threshold of 0.01. Information on each samples’ subgroup (cytogenetic MM subgroup, PC or “MM_OTHER” for MM samples with no cytogenetic information), condition (PC or MM) was not used by MOFA but was included for the analysis of the results. The variance explained was divided into 17 Latent Factors (LF).

The R script used to train the MOFA model can be found in:

MOFA/MOFA_top5k_var_peaks_and_genes_no_gender_no_TSS_train_model.R

The bash script to run the MOFA R script can be found in:

MOFA/MOFA_train_model_script.sh

Models using different random initializations were trained. Minimal differences were detected between different initialisations (data not shown). Models were fit using only RNA-seq data, or both RNA-seq and ATAC data.

#### Silhouette score for samples

For each model, each sample was assigned a label determined by its cytogenetic subgroup (including unknown cytogenetic samples, considered to be in the same group), the per sample silhouette score was obtained by calculating Euclidean distances between samples either using all LFs as dimensions or only LF1 to LF5. This was done using the “silhouette” function from the “cluster” R package version 2.0.6. The mean silhouette score for each subgroup and model is shown.

### Chromatin state analysis

The Chromatin State Segmentations (12 states) by ChromHMM for the 173 cell types for the GRCh38 genome available to date in The DeepBlue Epigenomic Data Server (Fernandez et al., 2016) were retrieved. Only cell types from the B-cell lineage were kept, in total, 19 samples.

For the chromatin state heatmap, the differentially accessible candidate enhancers significantly correlated with protein coding gene expression were divided into 200bp windows. Each 200bp enhancer region was intersected with the cell states table to obtain the chromatin state for each cell type in that region using Bedtools intersect. For each candidate enhancer, a “predominant” state was calculated (i.e. the state which accounts for the largest number of 200bp windows). 931 unique enhancers were filtered, keeping only those where a predominant state was found in 15/19 cell chromHMM samples, leaving 832 candidate enhancer regions corresponding to 5,912 200bp windows. The matrix of predominant states was hierarchically clustered using the Gower metric with the daisy function from the “cluster” R package using Ward.D2 linkage (where a candidate enhancer had no predominant state in a given cell line, this was considered missing data). This clustering of the predominant states was used to determine the order of regions, and therefore order the matrix of 200bp windows, leaving 200bp windows from the same region together, but full enhancer regions ordered according to the clustering.

The 832 potential enhancers with a predominant state were classified using their predominant states. States 7 (Genetic Enhancer High), 9 (Active Enhancer High), 10 (Distal Active Promoter 2kb High) and 12 (Active TSS High Signal H3K4me3 H3K27Ac) were classified as enhancer states. The final two were classified as enhancers as have removed annotated and unannotated TSS and it has recently been suggested that H3K4me1 is not a requirement for enhancer driven transcription in mESC(Dorighi et al., 2017) and in Drosophila melanogaster(Rickels et al., 2017) and that highly active enhancers are marked by H3K4me3 and not H3K4me1 in flies and mESCs(Henriques et al., 2018). Regions were classified on the following criteria:

Regions were first divided into 2 classes based on B-cell states (all cell types except PC, MM, MM cell lines) 12/19 cell types into:

- Category 1: Deactivated enhancers in B-cells: 3 or less out of 12 B cell types having an active enhancer state.
- Category 2: B-cell activated enhancers: B cell activated enhancers: 4+ B cell types having an active enhancer state.

Category 1 (deactivated in B-cells) was then subdivided into:

- De novo MM enhancers: 0/2 PC cells and 2+ (2 or more)/4 MM having an active enhancer state.
- All cell types deactivated regions/no predominant state: 0/2 PC cells and 1 or 0/4 MM.
- PC enhancers, deactivated in MM and Bcell: 1+/2 PC and 1 or less/4 MM.
- PC and MM only enhancers: 1+/2 PC and 2+/4 MM.

Category 2 (generally activated B cell) was subdivided into:

- MM only reactivated B cell enhancers: 0/2 PC cells and 2+/4 MM having an active enhancer state.
- PC and MM deactivated regions/no predominant state: 0/2 PC cells and 1 or 0/4 MM.
- Deactivated MM, PC and B cell enhancers: 1+/2 PC and 1 or 0/4 MM.
- PC, MM, B-cell enhancers: 1+/2 PC and 2+/4 MM.

For the analysis of CLL samples, representative samples for ATAC-seq and chromatin state were chosen for naïve, memory and germinal B-cells, tonsillar plasma cells and CLL samples, and ATAC and ChromHMM tracks downloaded from http://inb-cg.bsc.es/hcli/IDIBAPS_Biomedical_Epigenomics/CLL_Reference_Epigenome/. ATAC traces were created by averaging signal for each sample. To display chromatin state tracks, the emissions matrix from(Beekman et al., 2018) and Blueprint Epigenomics were manually inspected. The 12 states, although named differently matched almost exactly, and the states from (Beekman et al., 2018)were displayed using same colour as the colour used for the closest matching state from Blueprint.

### Inference of gene regulatory networks

Digital footprinting analysis was performed as previously described with minor modifications(Piper et al., 2015; Piper et al., 2013). Mapped reads files (bam files) were sub-sampled to the minimum depth across all samples and merged based on their corresponding cytogenetic subgroup (ND, MMSET, MAF, HD, CCND1). Consensus chromatin accessibility regions were obtained by merging open ATAC-seq peak files from individual samples for each subgroup. The wellington_footprints.py command from the pyDNAse/Wellington(Piper et al., 2013) package was used to obtain footprints on the consensus ATACseq regions for each subgroup, using the parameters: -fdr 0.05 -fdrlimit -10 -A. Predicted binding maps for a set of 769 highly curated human transcription factor motifs from the HOCOMOCOv1(Kulakovskiy et al., 2018) collection on identified footprints were generated for each subgroup using the annotatePeaks.pl script of the Homer_v4.9 package(Heinz et al., 2010). Transcription factor motif occurrences were also calculated for the consensus accessible chromatin regions for each subgroup and used as background. Only transcription factors with expression of TPM >=10 in at least one sample within each subgroup were considered for downstream analysis. In each subgroup, the relative adjusted frequency for each transcription factor t was calculated as:

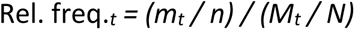

where,
*mt* is the number of motif occurrences in *n* total number of footprints

*Mt* is the number of motif occurrences in *N* total number of consensus ATAC-seq regions.

Footprint plots were generated with the dnase_average_profile.py command from the pyDNAse/Wellington package, using the ATACseq mode (-A). Transcription factor network visualization and analysis for each subgroup was performed using Cytoscape 3.5 4.

### Differential Footprinting on *CCND2*^*hi*^ versus *CCND2*^*lo*^ HD samples

Differential footprinting was performed in patients with HD myeloma with high (n=4) and low (n=4) *CCND2* mRNA expression (mean normalized count 0.23 vs 138.9, p= 0.016). Bam files were sub-sampled to the minimum depth across 8 samples and merged according to their *CCND2* levels (high, low). Consensus open chromatin regions were obtained by merging the ATAC-seq peak files from each sample. The wellington_bootstrap.py command from the pyDNAse/Wellington(Piper, Elze et al. 2013, Piper, Assi et al. 2015) package was used to obtain differential footprints on the consensus ATACseq regions for each subgroup, using the parameters: -fdr 0.05 - fdrlimit -8 -A. We considered only footprints with wellington score >10 for downstream analysis. Predicted differential binding maps for human transcription factor motifs from the HOCOMOCOv12 collection were obtained using the annotatePeaks.pl script of the Homer_v4.9 package(Heinz et al., 2010). For each TF with expression of TPM >=10 in at least one sample, the differential predicted frequency was calculated for each *CCND2* super-enhancer constituent region as:

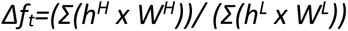

where,
*Δf*_*t*_ is the differential predicted frequency for transcription factor *t h* is the Homer motif purity score on differential footprints

*W* is the Wellington differential footprint probability score

The superscripts *H* and *L* refer to CCND2^High^ and CCND2^Low^ conditions, respectively.

### Data and code availability

Custom code used in this work is available at: https://github.com/sudlab/alvarez_et_al.

Raw sequencing data, peak sets and gene quantifications are deposited in GEO Accession GSEXXXXXX

