## Supplementary material for "Over-accessible chromatin links myeloma initiating genetic events to oncogenic transcriptomes and aberrant transcription factor regulatory networks": Supp Figures

**Supplementary Figures**

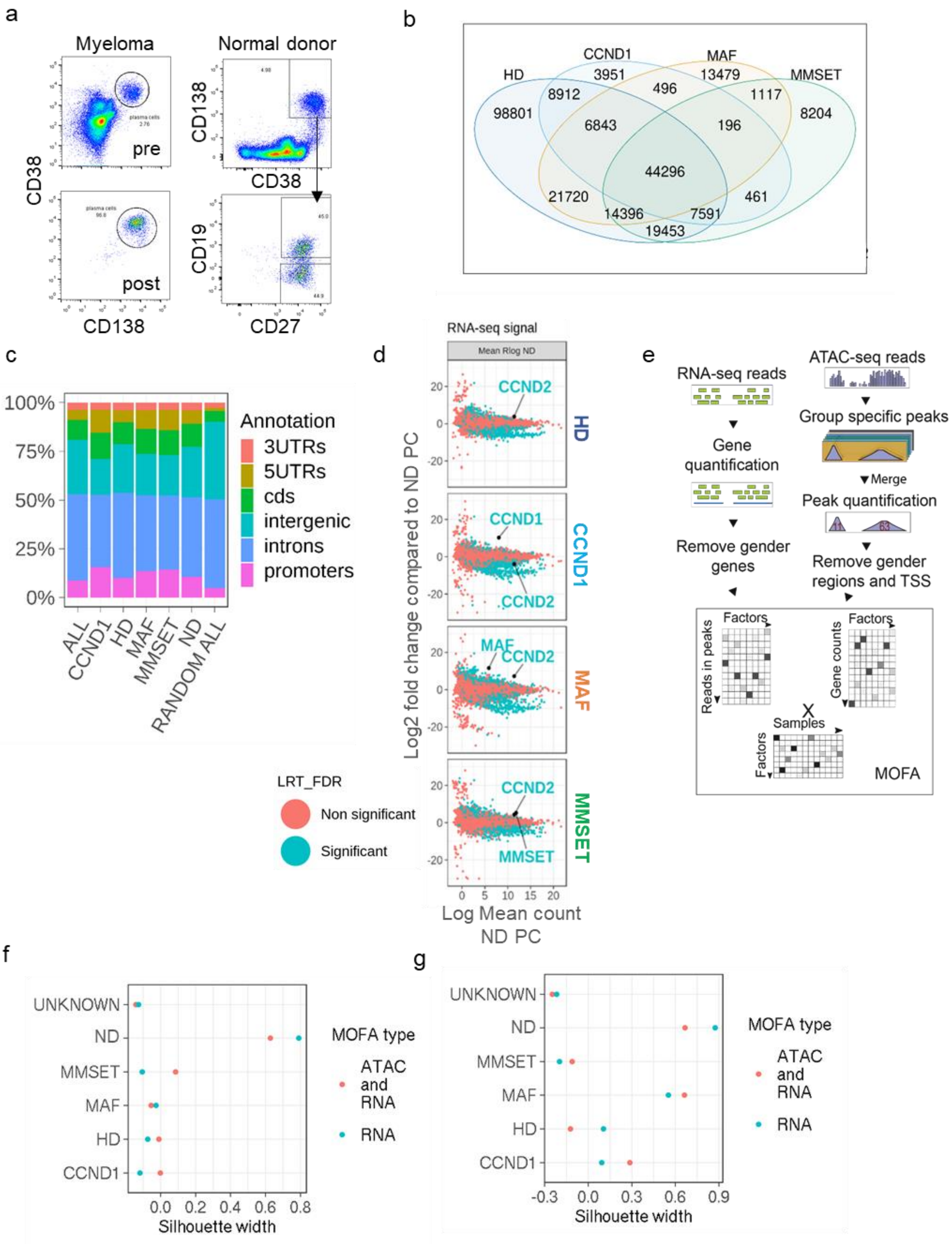


**Supplementary Figure 1 (related to Fig 1)**

**a)** Pre- and post-immunomagnetic bead selection of myeloma bone marrow PC (left); normal donor bone marrow PC were selected using flow-sorting according to indicated markers. In two samples both CD19$^{+}$ and CD19$^{-}$ PC were obtained while in a third sample only CD19$^{-}$ PC were obtained.

**b)** Venn diagram showing peaks discovered in ATAC-seq from MM samples with different genetic abnormalities.
**c)** Genomic annotation of peaks discovered in samples from each subtype. Each peak may overlap more than one context. 'Random All' refers to a set of size match regions per chromosome placed randomly.
**d)** MA plots correlating differential gene expression between subgroup myeloma PC and normal donor PC to mean gene expression level. Oncogenes of relevance for each subgroup are indicated.
**e)** Overview of the MOFA process**.**
**f & g)** Comparison of MOFA models built with both ATAC-seq and RNA-seq data, or RNA-seq data only. Performance in separating subtypes is measured as the silhouette score for members of that subtype compared to all other subtypes. High silhouette score means better clustering. Distance between samples is calculated as using Euclidean distance based on all (f) or the first 5 latent factors (g).

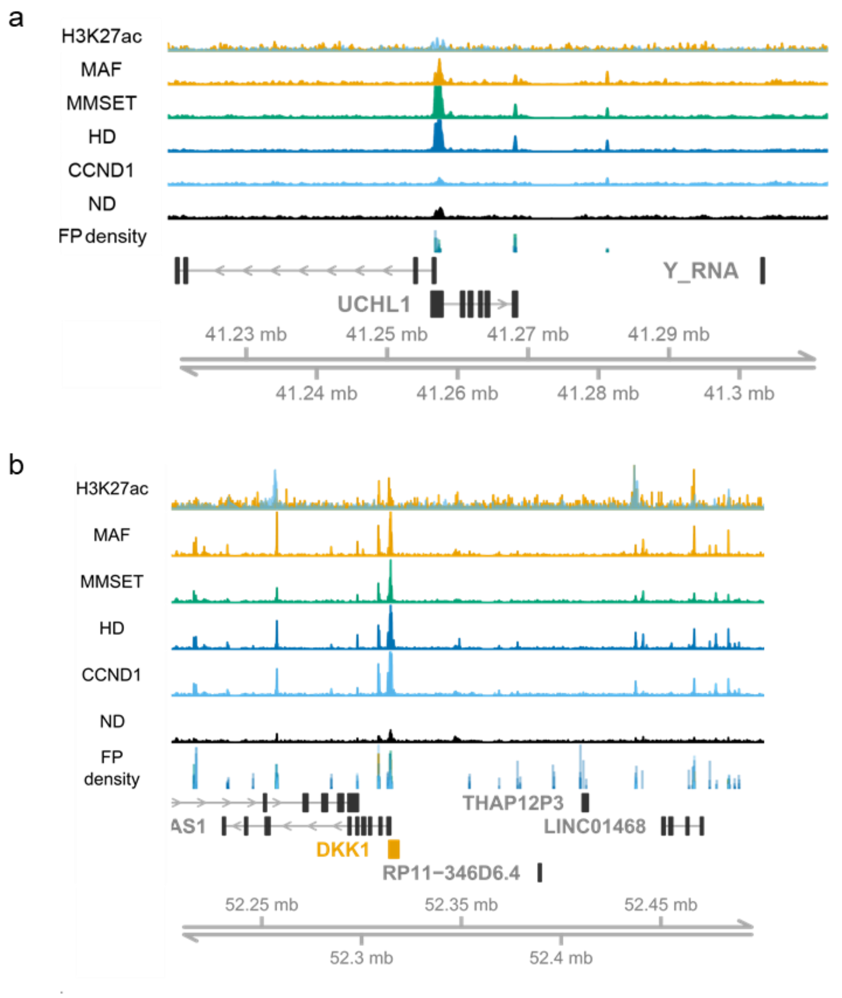


**Supplementary Figure 2 (Related to Fig 3)**
**a)** ATAC and H3K27ac signal around the *UCHL1* gene and **b)** the *DKK1* gene, both of which are upregulated in MM.
Tracks show normalised ATAC-seq (named after subtype), H3K27ac signal in *MAF* translocated cell line (Orange) and *CCND1* translocated cell line (blue) and density of footprints in the ATAC-seq signal (as called by Wellington).

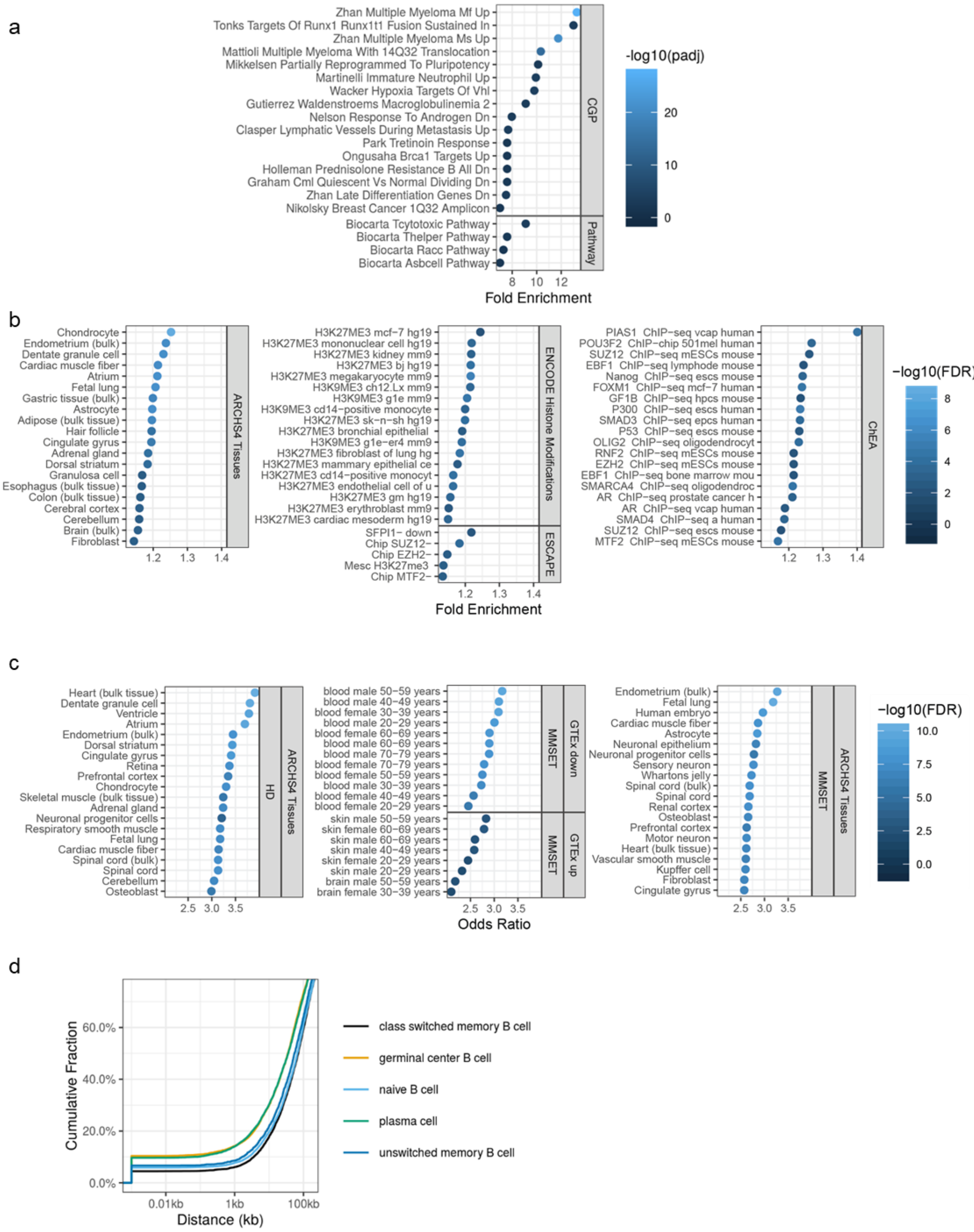


**Supplementary Figure 3**

Selected gene enrichment results of upregulated genes within 1Mb of a non-TSS ATAC peak of increased accessibility.

**a)** Upregulated genes within 1Mb of putative enhancers compared with all genes using the MSigDB curated pathways gene sets; CGP-Chemical/Genetic Perturbation, Pathway – Biocarta, KEGG, Reactome and Wikipathways.

**b)** The same genes compared to only upregulated genes using a selection of gene-set categories from Enrichr.

**c)** Genes upregulated in specific myeloma subgroups within 1Mb of putative enhancers from the same subgroup compared to upregulated genes using a selection of gene-set categories from Enrichr. Subgroup shown in the sidebar.

All enrichments BH corrected p-value < 0.01. A maximum of 20 gene sets is shown in any one category.

**d)** Distance of putative pan-MM enhancers from regions marked as Polycomb repressed chromatin in ChromHMM from B cell lineages.

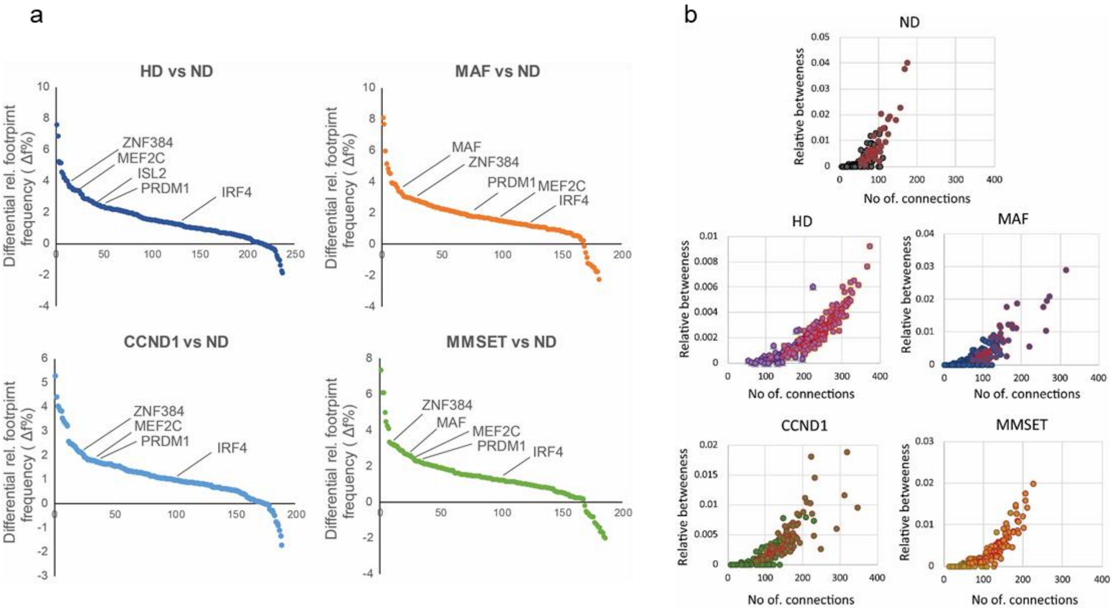


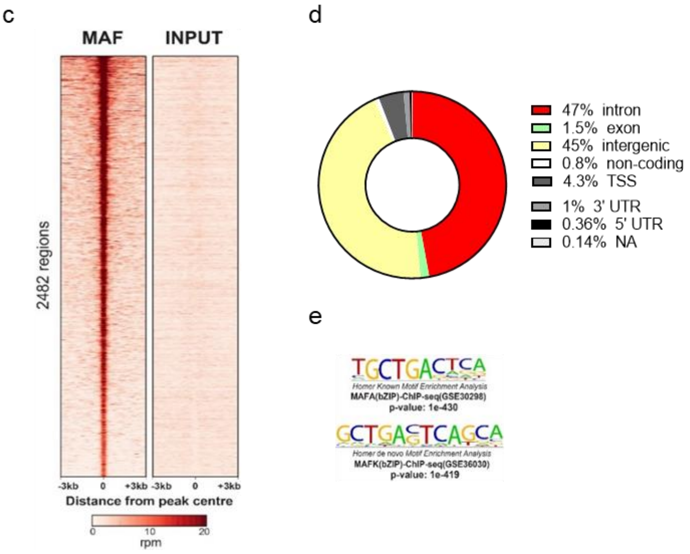


f

| Motifs | P value (-log$_{10}$) |
|---|---|
| MafA | 1150 |
| MafB | 1083 |
| MafF | 767 |
| MafK | 744 |
| IRF8 | 534 |
| Bach2 | 511 |
| Bach1 | 463 |
| IRF3 | 430 |
| IRF1 | 389 |
| NF-E2 | 358 |
| IRF2 | 343 |
| IRF4 | 341 |
| AP-1 | 329 |
| PU.1 | 315 |
| Fra2 | 315 |
| Nrf2 | 294 |
| Atf3 | 294 |
| Jun | 285 |
| Fosl2 | 279 |
| Fra1 | 270 |
| BATF | 269 |
| JunB | 265 |
| RUNX1 | 261 |
| ISRE | 258 |
| RUNX | 252 |
| RUNX2 | 229 |
| Fli1 | 215 |
| ETS1 | 199 |
| Etv2 | 187 |
| ERG | 170 |

**Supplementary Figure 4 (Related to Fig 4 & 5)**

**a)** Relative footprint frequency of indicated TF in myeloma subgroups and normal donor (ND) PC.

**b)** Biological networks analysis depicts the number of connections and the relative betweenness centrality of each active TF across MM subgroups and normal donor PC. TFs with active auto-regulatory loops are highlighted in red outline.

**c)** Heatmap illustration of MAF-bound genomic regions in MAF-translocated MM.1S cells, as compared to input control.

**d)** Genomic annotation of MAF cistrome in MM.1S cells.

**e)** Weblogo representation of MAF motifs found to be significantly enriched in MAF-bound regions in MM.1S cells.

**f)** TF motif discovery in MAF-bound genomic regions in MM.1S cells.

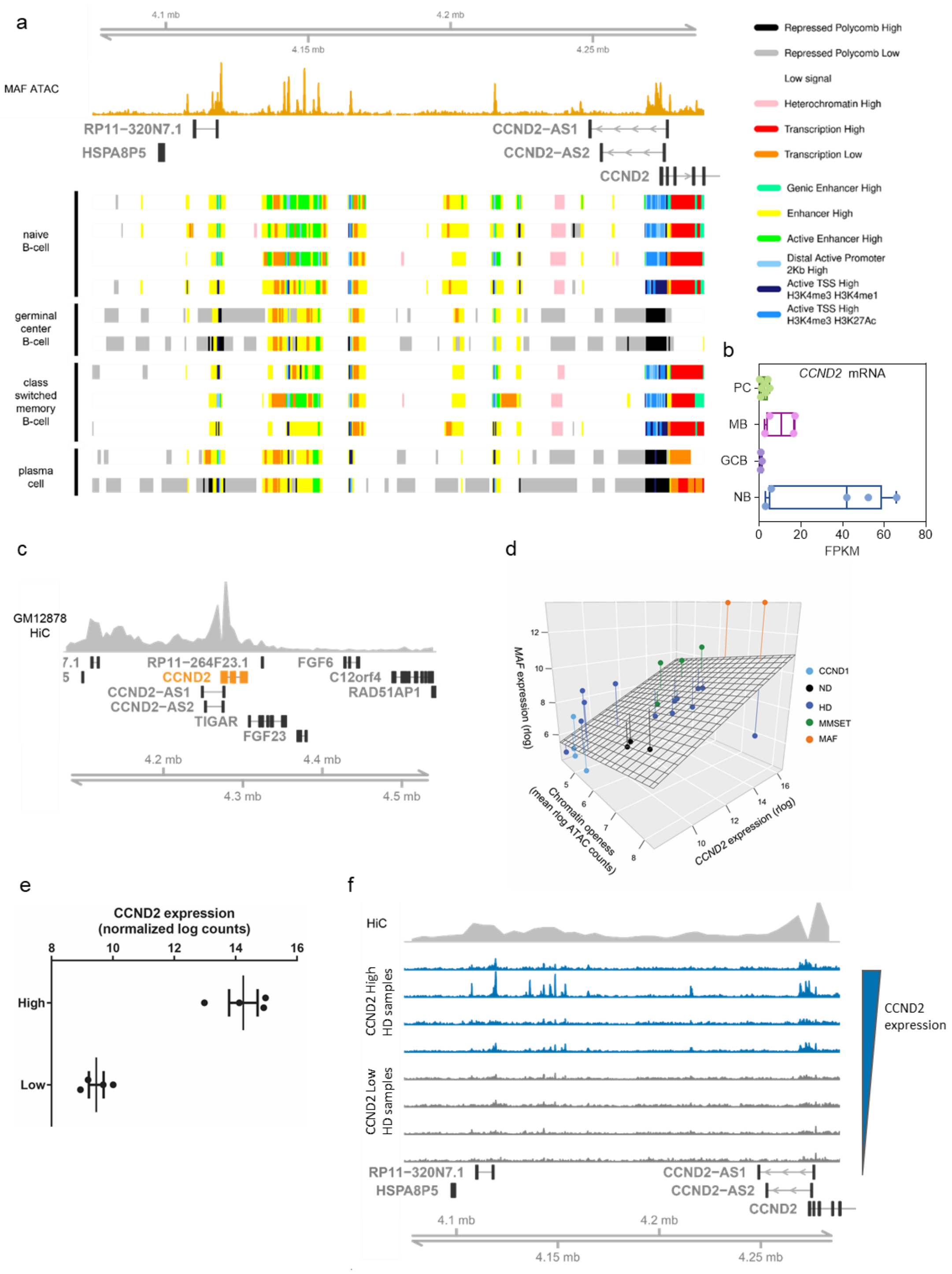


**Supplementary Figure 5 (related to Fig 6)**

**a)** The developmental origins of the *CCND2* super-enhancer as determined by ChromHMM state analysis across different B-cell types (naïve, germinal center, class-switched memory, plasma cells). ATAC-seq signal of MAF primary cells is also presented on the top panel**.**

**b)** *CCND2* expression in normal plasma cells (PC), memory B cells (MB), germinal center B cells (GCB) and naïve B cells (NB).

**c)** Hi-C signal for 3D genomic interactions around *CCND2* locus with its promoter region in GM12878 B cells.

**d)** 3D scatter plot displaying the correlation among chromatin accessibility of *CCND2* enhancer, *CCND2* and *MAF* gene expression. The distance of each point to the linear regression fitting surface is shown as a line from the point.
**e)** *CCND2* RNA-seq expression levels and
**f)** ATAC-seq signal tracks in *CCND2*$^{high}$ and *CCND2*$^{low}$ patient samples within the HD molecular subgroup.

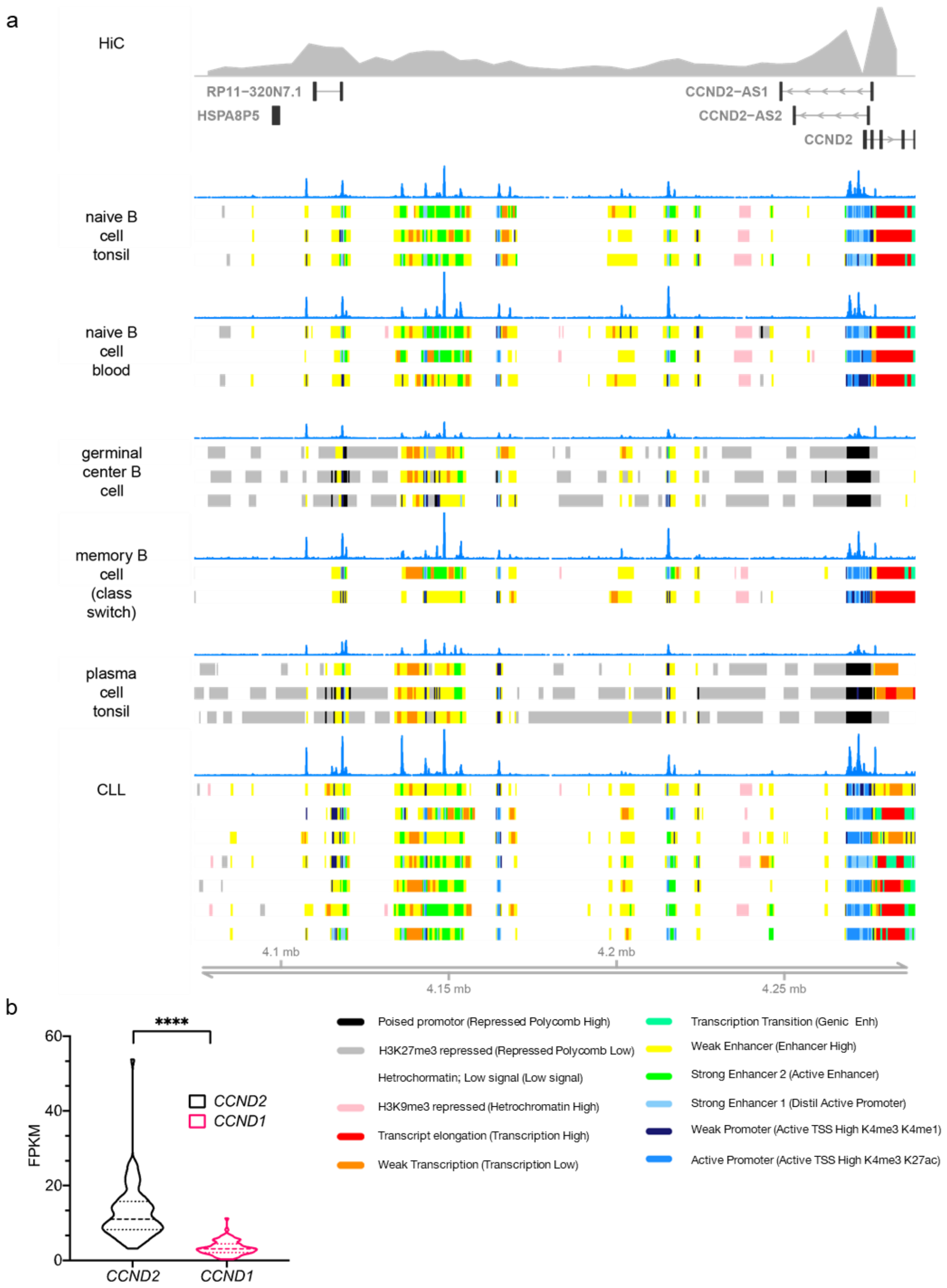


**Supplementary Figure 6 (related to Fig 6)**
**a)** Chromatin landscape of the *CCND2* locus in chronic lymphocytic leukemia (CLL) and associated normal B cell types from Beekman *et al*. Colour tracks show ChromHMM states. Key shows Beekman *et al* annotation of chromatin state, with the closest matching state from

the model used by Blueprint epigenomics (as shown in Figure 2 and Supp Figure 5). Traces are average ATAC-seq signal over samples of that type. Hi-C track shows Hi-C signal associated with the *CCND2* promoter from GM12878 B cells.

**b)** RNA-seq signal for *CCND2* and *CCND1* from CLL samples (n=78).
